# Failure to mate enhances investment in behaviors that may promote mating reward and impairs the ability to cope with stressors via a subpopulation of Neuropeptide F receptor neurons

**DOI:** 10.1101/2021.04.27.441612

**Authors:** Julia Ryvkin, Liora Omesi, Yong-Kyu Kim, Mali Levi, Hadar Pozeilov, Lital Barak-Buchris, Bella Agranovich, Ifat Abramovich, Eyal Gottlieb, Avi Jacob, Dick R. Nässel, Ulrike Heberlein, Galit Shohat-Ophir

## Abstract

Living in dynamic environments such as the social domain, where interaction with others determines the reproductive success of individuals, requires the ability to recognize opportunities to obtain natural rewards and cope with challenges that are associated with achieving them. As such, actions that promote survival and reproduction are reinforced by the brain reward system, whereas coping with the challenges associated with obtaining these rewards is mediated by stress-response pathways, the activation of which can impair health and shorten lifespan. While much research has been devoted to understanding mechanisms underlying the way by which natural rewards are processed by the reward system, less attention has been given to the consequences of failure to obtain a desirable reward. As a model system to study the impact of failure to obtain a natural reward, we used the well-established courtship suppression paradigm in *Drosophila melanogaster* as means to induce repeated failures to obtain sexual reward in male flies. We discovered that beyond the known reduction in courtship actions caused by interaction with non-receptive females, repeated failures to mate induce a stress response characterized by persistent motivation to obtain the sexual reward, reduced male-male social interaction, and enhanced aggression. This frustrative-like state caused by the conflict between high motivation to obtain sexual reward and the inability to fulfill their mating drive impairs the capacity of rejected males to tolerate stressors such as starvation and oxidative stress. We further show that sensitivity to starvation and enhanced social arousal is mediated by the disinhibition of a small population of neurons that express receptors for the fly homologue of neuropeptide Y. Our findings demonstrate for the first time the existence of social stress in flies and offers a framework to study mechanisms underlying the crosstalk between reward, stress, and reproduction in a simple nervous system that is highly amenable to genetic manipulation.

## Introduction

Living in a social environment involves diverse types of interactions between members of the same species, the outcomes of which affect health, survival, and reproductive success^1^. Coping with the challenges and opportunities associated with this dynamic environment requires individuals to rapidly process multiple sensory inputs, integrate this information with their own internal state and respond appropriately to various social encounters^1–7^. Encounters that hold opportunities to secure resources, mating partners, and a higher social status are considered rewarding and are, therefore, reinforced by the brain reward systems, whereas the failure to obtain such rewards due to high competition, lack of competence, or aggressive encounters with competitors are perceived as stressors^8–10^.

It has been appreciated for quite some time that stress-response mechanisms allow animals to cope successfully with challenges encountered when living in social groups^8, 11–14^. For instance, high levels of stress hormones promote attentive care during maternal behavior in humans^15^, and the CRF-Receptor2 and its ligand Urocortin-3 are necessary for coping with social engagement in mice^16–19^. Moreover, high levels of CRF are correlated with increased motivation to obtain natural rewards, as documented in individual cichlid fish that ascend in the hierarchy of their social group^20^. While these are examples of the ways by which stress response mechanisms improve the ability of individuals to cope with social challenges, some types of social stress can induce social defeat, increase drug consumption^21–26^, impair health and shorten lifespan^27–31^.

There is considerable anatomical overlap between brain regions that process reward and stress stimuli, as well as opposing functionality: exposure to natural rewards buffers the effect of stressors, while stressors such as social defeat can alter sensitivity to reward and increase the rewarding value of certain addictive drugs^27, 32–38^. An example of such opposing functions is seen in rodents where corticotropin-releasing factor (CRF) increases alcohol intake, and the binding of Neuropeptide Y (NPY) to NPY receptor Y1 on CRF-positive neurons within the bed nucleus of the stria terminalis (BNST) inhibits binge alcohol drinking^34, 39, 40^.

Similar responses to social stress and reward-seeking behaviors can be seen in a variety of animals, suggesting that the central systems facilitating survival and reproduction originated early in evolution and that similar ancient basic building blocks mediate these processes^41, 42^. In agreement with this concept, we and others showed that *Drosophila melanogaster* can adjust its behavior and physiology to various social conditions^43–55^ and that the brains of mammals and fruit flies share similar principles in encoding stress and reward^43, 56–58^. For example, the fly homologue of the NPY signaling system (i.e., NPY and its receptor) functions in processing natural and drug rewards, decreases aggressive behaviors and suppresses responses to aversive stimuli such as harsh physical environments^43, 59–69^. Moreover, similar to the essential role that NPY\Y1 signaling plays in various processes affecting health and lifespan in mammals^70–74^, activation of neuropeptide F (NPF) neurons in male flies leads to decreased resistance to starvation and decreased lifespan^75, 76^.

We previously showed that successful mating, and, more specifically, ejaculation, is rewarding for male fruit flies and reduces the motivation to consume drug reward^43, 77, 78^. While most research focuses on mechanisms that encode sexual reward^43, 60, 75, 77, 79–84^, it remains unknown whether failure to obtain sexual reward is simply a lack of reward or is perceived as a stressor. Here we used the courtship suppression paradigm in *Drosophila*^45, 85–87^ to model the effects of failures to obtain sexual reward on different aspects of male behavior. We discovered that failure to mate induces a stress-like response characterized by a larger investment in actions that can enhance the odds of obtaining sexual reward while, at the same time, reduces the ability of rejected males to endure starvation and oxidative stressors via the disinhibition of a small subset of NPF-receptor neurons.

## Results

### Failure to mate promotes social avoidance

To explore the way by which failure to obtain rewards affects behavior, and to ask whether failure to obtain reward is simply a lack of reward or is perceived as a challenge, we used the courtship suppression paradigm to induce repeated events of sexual rejection and assayed various aspects of male behavior. As a starting point, we employed the agnostic nature of the FlyBowl system to compare multiple behavioral features of male flies that experienced repeated failures to mate over the course of 4 days (“rejected”), to that of controls that experienced successful mating (“mated”), or lack of any sexual and social experience (“naïve-single”). Following the experience phase, 10 flies of each cohort were placed in circular arenas, and their behavior was recorded for 30 min and analyzed using the FlyBowl suite of tracking and behavior analysis softwares^88–90^ (Fig. 1A). The tracking data obtained was used to calculate various kinetic features, including velocities, distances and angles between flies and their relative differences across time, and to train 8 types of behavior classifiers using the Janelia Automatic Animal Behavior Annotator (JAABA) (Fig. 1B, S1)^88^.

**Figure 1.**
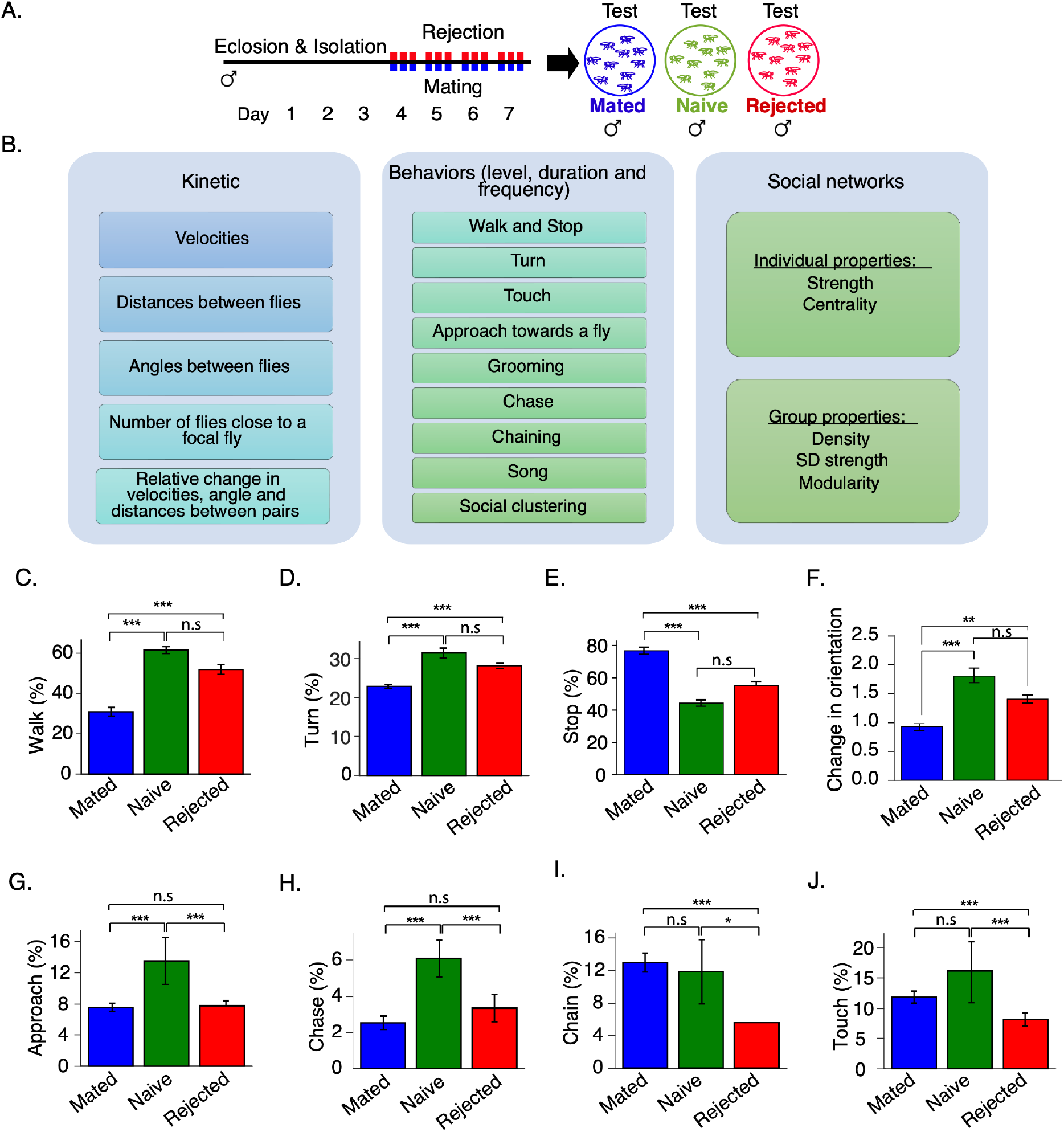
Failure to mate modifies action selection upon encounters with rival male flies. **A.** Schematic representation of sexual experience and behavioral analysis. Rejected males experienced 1h long interaction with non-receptive female flies, 3 times a day for 4 days. Mated males experienced 1h long interactions with a receptive virgin female, 3 times a day for 4 days. Male flies with no sexual experience are kept in social isolation (naïve-single). The social group interaction of the 3 cohorts is recorded for 30 min using the FlyBowl system. **B.** List of kinetic features, 8 behaviors and network parameters produced using the FlyBowl system. **C-J** Average percentage of time mated (blue) naive (green) and rejected (red) took to perform walk (C), turn (D), stop (E), change orientation (F), approach (G), Chase (H), chain (I) and touch (J) behaviors. n = 15, 10, 15 for mated, naïve-single and rejected respectively. one-way ANOVA followed by Tukey’s and FDR correction for multiple tests *p < 0.05, **p < 0.01, ***p < 0.001. Error bars signify SEM.

Examining features that reflect activity levels, such as time spent walking, number of turns, time spent stopping, and average speed, revealed similar activity levels in rejected and naïve-single males, which were significantly higher than those in mated males (Fig. 1 C-F). Analysis of specific social behaviors indicated an overall reduction in the degree of social interaction in rejected males than in naïve-single males (Fig. 1 G-J). This included a lower number of approaches towards other males (Fig. 1G), low levels of male-male chase behavior (Fig. 1H), reduced formation of chains composed of multiple chasing males (Fig. 1I), and fewer close touch encounters (Fig. 1J). Taken together, the results indicate that while rejected male flies exhibit higher activity levels that are similar in their extent to that of naïve-single males, they have overall reduced levels of behaviors associated with social interaction.

To extend the analysis of their social behavior, we next investigated the network structures of the various fly cohorts, by analyzing features that describe the properties of their respective social networks and the individuals within them, including the density of the network and the strength of individuals within it (Fig. 2A)^88^. Interaction events were defined using the following parameters: focal fly can see the other fly (requirement: at least 2 seconds in which the focal fly was at a distance of >8mm from the other fly and in which the visual field of view of the focal fly was occupied by the other fly (angle subtended >0); and network weights, i.e., the overall duration of interactions (emphasizing long-lasting interactions) or overall number of interactions (emphasizing short interactions) between each pair of flies (Fig. 2B-E). Analysis by duration indicated that the social networks of rejected males are less dense (Fig. 2B) and that the average strength of individual flies in these networks is lower compared to mated and naive male flies (Fig. 2C). Likewise, analyzing network features by number of interactions identified reduced density and low strength of the individuals than naive males (Fig. 2 D, E). These findings suggest that rejection promotes the formation of sparser groups containing individuals with reduced social interaction.

**Figure 2.**
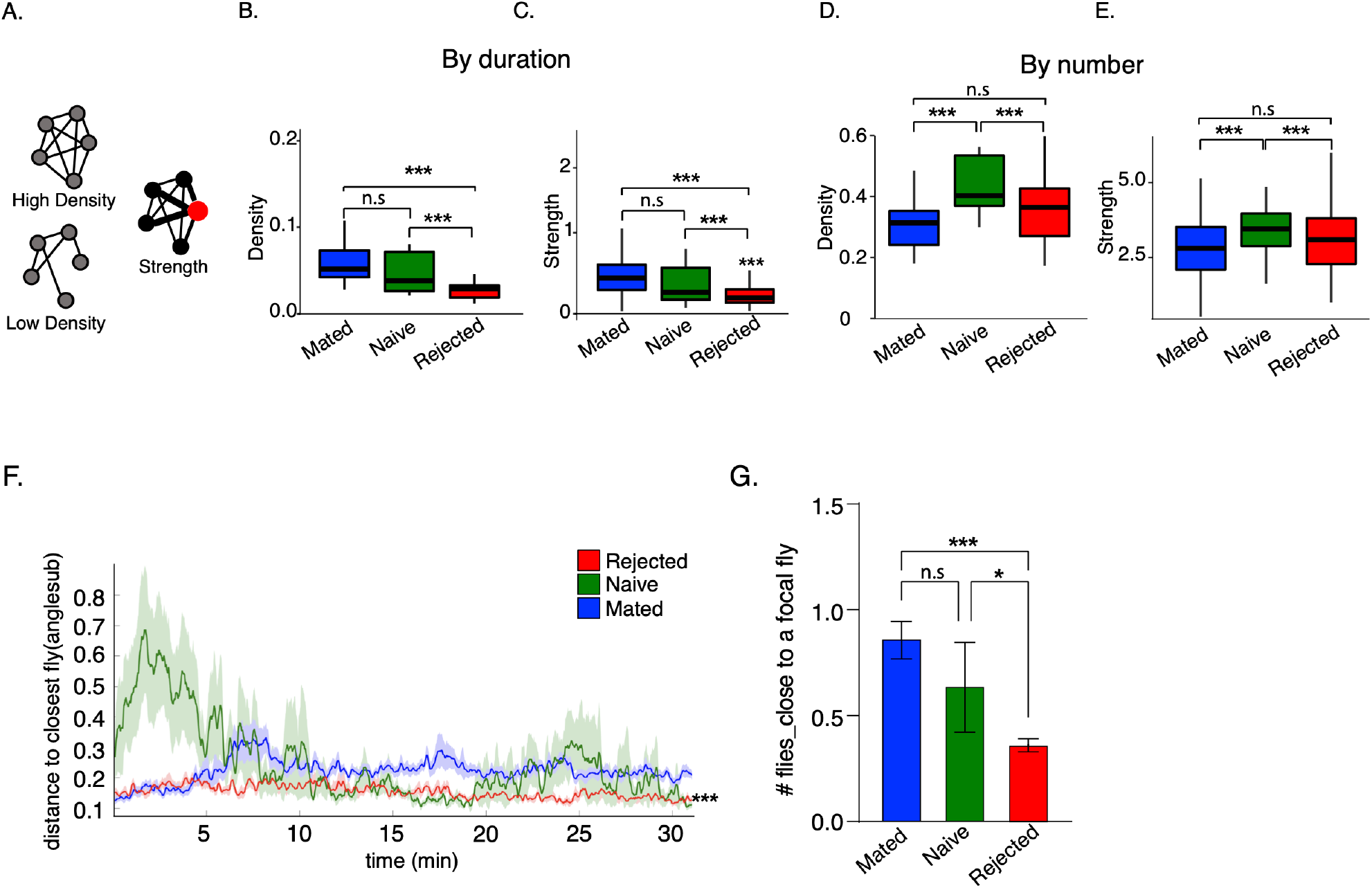
Sexual rejection promotes social avoidance. **A.** Illustration of network parameters. Density of networks represents how saturated they are compared to the maximum possible. Strength is proportional to vertex size (high in red individual). **B-E.** Social network analysis of groups composed of mated (blue) naïve-single (green) and rejected (red). Network density, and strength calculated by network weights according to duration (B-C) or number of interactions (D-E). Kruskal–Wallis test followed by Wilcoxon signed-rank test and FDR correction for multiple tests *p < 0.05, **p < 0.01, ***p < 0.001, Box plot with median IQR. **F.** Rejected male flies maintain large distances between flies along (anglesub), statistical significance was tested on averages across time. one-way ANOVA followed by Tukey’s and FDR correction for multiple tests *p < 0.05, **p < 0.01, ***p < 0.001. Error bars signify SEM. **G.** Average number of flies close to a focal fly. n = 15, 10, 15 for mated, naïve and rejected respectively. one-way ANOVA followed by Tukey’s and FDR correction for multiple tests *p < 0.05, **p < 0.01, ***p < 0.001. Error bars signify SEM.

The reduced network density of rejected male flies prompted us to compare the average distance between individuals of each group along the experiment, as indicated by the average field of view occluded by another fly (anglesub), a feature that increases as the distance between individuals decreases (Fig. 2F). This analysis revealed that rejected male flies maintained significantly low values of anglesub, suggesting that rejected males maintain long distances between one another across the experiment. This finding is also supported by a reduced number of flies near a focal rejected fly compared to mated and naive male flies (Fig. 2G). Given that rejected male flies exhibit similar activity levels as naive male flies, and presumably a higher probability to encounter other flies, their reduced network density and high inter-individual distance suggest that rejected individuals actively avoid social interactions with other flies, resulting in low-density groups.

The behavioral signatures of the three cohorts across all 60 parameters (kinetic features, velocities, distances and angles between flies, scores for 8 behaviors, their frequencies as well as network features), are summarized in a scatter plot of normalized differences and are divided into 4 main categories: activity, interaction, coordination between individuals and social clustering, i.e., the aggregation of males to form clusters (Supp Fig.2). The existing similarities and differences between the three cohorts indicate that sexual rejection induces a discrete behavioral state that differs from successful mating and lack of mating and point to sexual rejection as the major contributor to reduced social interaction.

### Failure to mate promotes stress responses that can increase the odds of obtaining a sexual reward

The similarity between some of the behavioral features exhibited by sexually rejected male flies to stress responses in other animals, including social avoidance, enhanced activity/arousal, and increased consumption of drug rewards^43^, suggests that failure to mate induces a stress response in male flies. This prompted us to investigate whether failure to mate induces the loser-like state observed in socially defeated animals^16, 91^ or, rather, a high motivation to obtain a sexual reward, as described in several species upon the omission of an expected reward^92–102^. To discriminate between these two options, we analyzed the behavioral responses of rejected males in aggressive and mating encounters. If rejection promotes a loser-like state, rejected males are expected to exhibit low levels of aggression towards other male flies and reduced investment in mating. Conversely, if it promotes a high motivational state, rejected males should display enhanced aggression and higher investment in mating-related behaviors.

We compared the aggression levels of rejected and mated male flies, as both cohorts experienced interaction with females, unlike naïve-single flies, which were socially isolated, a condition known by itself to promote aggression. We discovered that when paired together, rejected male flies exhibited significantly higher displays of aggression in comparison to pairs of mated male flies (Fig. 3A). In mixed pairs, the rejected male exhibited a far greater number of lunges compared to its mated opponent (Fig. 3B, C), indicating that failure to mate does not induce a loser-like state. To extend this analysis to mating-related actions, we compared the mating duration of rejected males to that of control naïve/virgin male flies, as the investment in mating of the mated cohort is shaped by their previous mating events. When allowed to mate with virgin female flies, the copulation duration of rejected male flies was 25% longer (3.5 minutes longer) than the control males (Fig. 3D). This was accompanied by an increase in the expression levels of genes associated with reproductive success compared to control males, including a two-fold increase in the transcript levels of *Sex-Peptide (Acp70A)* and *Acp-63,* both of which facilitate post-mating responses and fertility in female flies (Fig. 3E)^103^. Taken together, these findings suggest that repeated events of rejection promote a high motivational state rather than a loser-like state.

**Figure 3.**
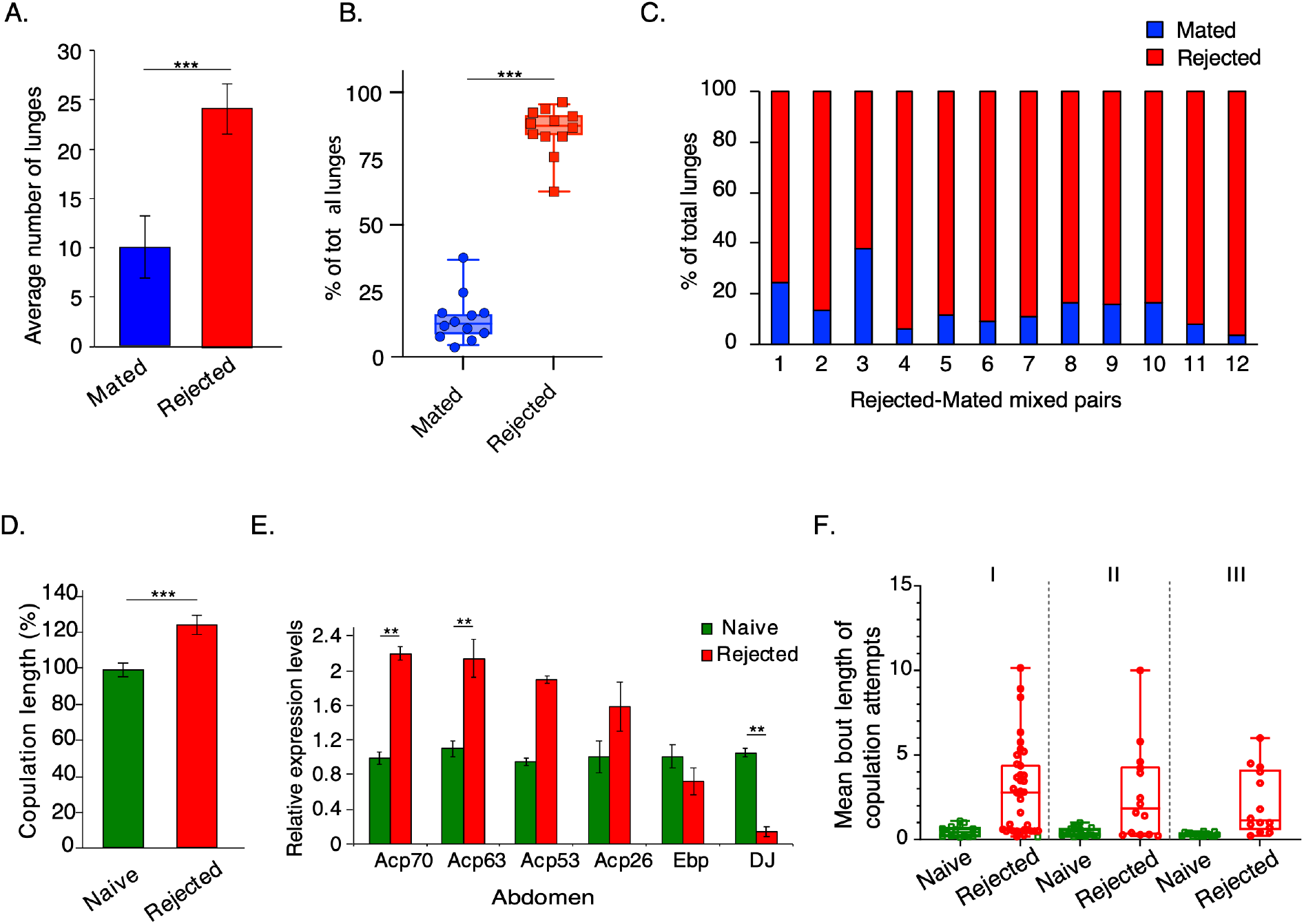
Sexual rejection modulates competitive behaviors and ejaculate composition. **A.-C.** Aggression display (number of lunges) was compared between pairs of rejected and mated male flies (n=16, statistical significance determined by T-test, *p*< 0.005 **(A),** and mixed pairs (n=12) **(B-C).** The log2 ratio between the number of lunges in rejected and mated flies was calculated for each pair, and then a one-sample T-test was performed to test whether the mean ratio was significantly different than 0, *p*<0.005. Data is presented as the mean ± SEM. **D.** Duration of copulation in rejected vs. naïve-single male flies. Statistical significance was determined by T-test, *p* < 0.001. Data is presented as mean ± SEM, n=25. **E.** Relative transcript levels of candidate genes expressed in abdomens of rejected and naïve-single male flies were quantified by qRT-PCR, n = 6 independent experiments of 15–20 fly heads and abdomen. Statistical significance was determined by Student’s T-test with Bonferroni correction for multiple comparisons. **, *p* <0.01 ***, *p* < 0.005. **F.** Virgin male flies were exposed to either mated or virgin females for three 1h sessions, and their behavior was recorded. At the end of each session, the females and males from the control cohort (Naive) were removed, and the males that experienced rejection were kept isolated in narrow glass vials for 1h. Mean bout duration of their copulation attempts was measured in the first (I), second (II), and third (III) sessions. Student’s t-test or Mann-Whitney was performed with FDR correction for multiple comparisons. \**p*<0.05, \*\*\**p*<0.001. n = 41

To further explore the effects of rejection on mating drive, we performed a detailed analysis of the action selection exhibited by males towards mated females during the first day of the training phase that was used to generate the rejected cohort (three one-hour training interactions spaced by one-hour resting intervals). As controls we used in each training session virgin males that interacted with virgin females (Supp Fig.3A). The behavior of both cohorts during the first 10 min of each session was manually analyzed. In line with previous studies, the rejected cohort displayed marked courtship suppression, reflected by a reduction in the overall time spent courting in all 3 sessions (Supp Fig. 3B)^45, 85, 104^. Yet, the overall number of males exhibiting courtship action did not decline over the course of two days (Supp Fig. 3C), and interestingly, other aspects of their courtship, such as the overall number of licking actions and number of copulation attempts, were no less vigorous (Supp Fig. 3B). Notably, although the rejected males depicted longer latency to court mated females during the first encounter (consistent with an innate aversion to the male pheromone cVA^104, 105^), they overcame this aversion in subsequent sessions and initiated courtship at the same time as males that courted virgin females (Supp Fig. 3B). Another feature that may reflect their surprising persistence is the duration of copulation attempts, which was 6 times longer compared to the controls (Fig. 3F). This finding is suggestive of a conflict between high motivation to mate and repeated inability to fulfill this mating drive. Taken together, the behavioral responses of rejected males towards male and female flies imply that repeated failures to mate induce a “frustration-like” state where rejected males experience a conflict between their high motivation and inability to fulfill their mating drive.

### Sexual rejection increases sensitivity to stressors

Next, we assessed whether there is a cost for this “frustration-like” state by testing the ability of rejected males to endure starvation and oxidative stress. Rejected, mated and naïve-single males were conditioned over the course of two days, and each cohort was subsequently exposed to starvation (1% agarose) or oxidative stress (20mM paraquat), and the rate of survival over time was documented (Fig. 4A). If deprivation of sexual reward is a stressor, rejected male flies should be more sensitive to other stressors. Indeed, rejected males exhibited higher sensitivity to both starvation and oxidative stress compared to controls (Fig. 4 B,C). While the survival of mated and naïve-single cohorts declined to 50% after 22– 24h of starvation, in the rejected male flies this was reached after less than 18h (Fig. 4 B). Exposure to paraquat led to a 50% decline in survival after more than 17h for both the mated and naïve-single cohorts vs. less than 14 hours for rejected males (Fig. 4C). This demonstrates that sexual rejection promotes sensitivity to starvation and oxidative stress, further suggesting that multiple rejection events serve as a stressor, compromising the ability of male flies to cope with other stressors. Importantly, complete deprivation of both social and sexual interactions (i.e., naïve-single cohort) did not result in the same effect as active rejection, suggesting that the deprivation of an expected sexual reward, but not the lack of mating, increases sensitivity to additional stressors. The overall lifespan of rejected males was not affected by sexual deprivation (Supp Fig. 4A).

**Figure 4.**
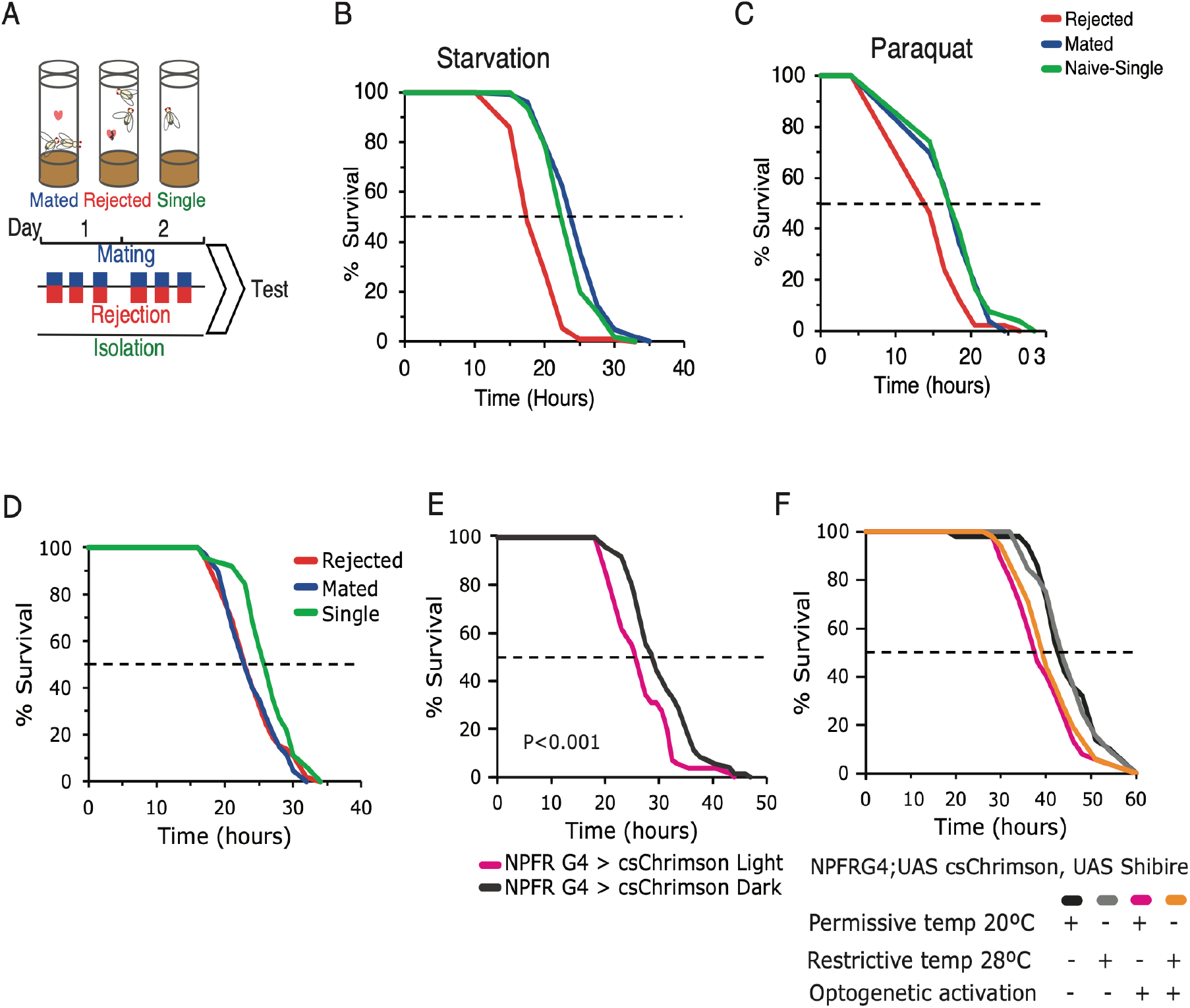
Repeated sexual rejection events increase sensitivity to stressors via NPFR neurons. **A.** Schematic representation of courtship conditioning: control males were introduced to either virgin, sexually receptive or sexually non-receptive females. As a result, males were either mated or rejected. The third cohort consisted of control single housed males that did not experience any social or sexual event (naïve-single). Encounters with females were repeated 3x a day for two days. **B**. Starvation resistance assay: rejected males (blue, n=91) compared to mated (red, n=102) and single housed (yellow, n=110) males, \*\*\**p*< 2E-16; mated vs single males, \**p*<0.05. **C.** Resistance to oxidative stress (20mM Paraquat): rejected males (blue, n=50) compared to mated (red, n=53) *p*<0.01** and naïve-single (yellow, n=54) males \*\*\**p*<0.001. Pairwise log-rank with FDR correction for multiple comparisons was performed in E,F. **D. .** Starvation resistance assayed on NPFR>RNAi mated (red, n=68), rejected (blue, n=68) and naïve-single (yellow, n=63) male flies. \**p*<0.05 rejected vs naïve-single, \*\**p*<0.01 mated vs naïve-single. Pairwise log-rank with FDR correction **E.** Activation of NPFR neurons promotes sensitivity to starvation. NPFR>csChrimson activation three times a day for two days (pink), NPFR>csChrimson w/o activation (grey) serve as controls. Starvation resistance of experimental (n=55) and control flies (n=72) was assayed. Log-rank test was performed, \*\*\**p*< 0.001. **F.** Male flies expressing UAS-csChrimson and UAS-Shibire^ts^ in NPFR neurons were subjected to three 5 min long optogenetic activations for three days, and their synaptic signaling was blocked at 28-29°C (light+ heat, orange, n=52). Positive control males (light+cold, pink, n=52), synaptic release block control (dark+heat, light gray, n=52), negative control (dark+cold, dark gray, n=50). Experimental and positive control flies showed no significant difference in resistance to starvation (*p*>0.05). Both experimental and positive control flies were significantly more sensitive to starvation than ‘dark+heat’, and ‘dark+cold’ flies (\*\**p*<0.01). Pairwise log-rank test with FDR correction for multiple comparisons was performed.

The sensitivity of rejected males to starvation was not due to the increased activity of the rejected cohort, as there was no difference in the circadian activity and sleep patterns between rejected males and the other two cohorts (Supp Fig. 4B-D). It also implies that the enhanced activity in the FlyBowl experiments does not reflect an inherent increase in the activity of individual flies, but rather an emergent property of their response to the presence of other males. Additionally, no difference was documented in body triglycerides (TAGs), glucose levels, or body weight between the cohorts (Supp Fig. 5A-D), suggesting that rejected male flies do not suffer from an energetic deficit that increases their sensitivity to starvation. Although targeted metabolite analysis of head tissues using liquid chromatography–mass spectrometry (LC-MS) revealed a unique profile for each cohort (Supp Fig. 5E), we did not document any significant accumulation or depletion of metabolites such as oxidative agents or in antioxidant activity and glucose levels. Thus, sexual rejection decreases the ability of males to cope with additional stressors, but this is not simply explained by metabolic deficits.

### Disinhibition of NPFR neurons increases sensitivity to starvation in male flies

The sensitivity of rejected males to acute stressors implies the existence of a link between the inability to obtain rewards and the stress response. Given the causal link between sexual rejection, NPF/R signaling and ethanol consumption in flies^43, 63^, and the role of the mammalian NPY-in mediating the crosstalk between reward and stress by inhibiting downstream neurons^34^, we postulated that rejection leads to disinhibition of NPFR neurons, which in turn sensitizes males to starvation. To test this hypothesis, we knocked down NPF-receptor in NPFR neurons and compared the sensitivity of naïve-single, rejected, and mated male flies to starvation stress. If sexual rejection reduces NPF signaling, which in turn disinhibits NPFR neurons, mated male flies should exhibit similar responses as rejected male flies. Our findings show that this manipulation abrogated the differences between mated and rejected cohorts (Fig. 4D), implying that the release of NPF and its binding to its receptor is necessary for the resistance of mated males to starvation stress. Moreover, mimicking rejection state by artificial activation of NPFR neurons in naïve-single males significantly increased their sensitivity to starvation compared to control flies (Fig. 4E), which further demonstrates that the disinhibition of NPFR in rejected males induces sensitivity to starvation stress. Interestingly, neurotransmitter release may not be required for starvation sensitivity triggered by the activation of NPFR neurons, as the activation of NPFR neurons and simultaneous inhibition of their synaptic transmission (using temperature sensitive Shibire), yielded similar levels of sensitivity to starvation as that of activation of NPFR neurons alone (Fig. 4F). These results suggest that sexual-deprivation-induced sensitivity to starvation may involve neuropeptide based signaling rather than dynamin-based neurotransmitter release.

To assess whether the activation of NPFR neurons can also mediates other behavioral phenotypes observed in rejected males, we assayed the behavior of NPFR>csChrimson flies during optogenetic activation in FlyBowl arenas. Interestingly, we observed minor difference in the behavior of the experimental flies compared to genetic controls (Supp Fig. S6A), indicating that disinhibition of all NPFR neurons is sufficient to increase sensitivity to starvation, but not to induce changes in male-male social interactions.

### Activation of a small subpopulation of NPFR neurons increases sensitivity to starvation and promotes social arousal

We next sought to determine which subset of the 100 NPFR neurons is responsible for the enhanced sensitivity to starvation. Given the possibility that sensitivity to starvation is mediated via neuropeptide release, we focused our attention to neuropeptide expressing neurons with shared expression with NPFR such as NPF^106^, Dh44^61^, and Tachykinin^107^. Activation of NPFR^NPF^ mutual cells (P1 and L1-l neurons) did not affect either the sensitivity to starvation or male-male social interactions (Supp. Fig. 6B, C), suggesting that NPF-expressing NPFR neurons, which presumably have the capacity for autoinhibition, are not responsible for these effects. Next, we tested Dh44 expressing neurons, as we previously documented similarity between DH44 and NPFR transcriptional programs^61^. We first validated the overlap between the two populations and discovered that all six pars intercerebralis (PI) Dh44 neurons are NPFR neurons as well (Fig. 5A). Activating Dh44 neurons did not affect sensitivity to starvation, nor did it lead to apparent effects on male-male behavioral responses (Fig. 5B, Supp. Fig 6D). In addition, knocking down the expression of Dh44 in NPFR neurons did not change the sensitivity to starvation (Supp. Fig 6E). Altogether, these findings suggest that although DH44 neurons are the functional homologue of mammalian CRF, Dh44 signaling does not mediate the sexual reward deprivation stress response in flies.

**Figure 5.**
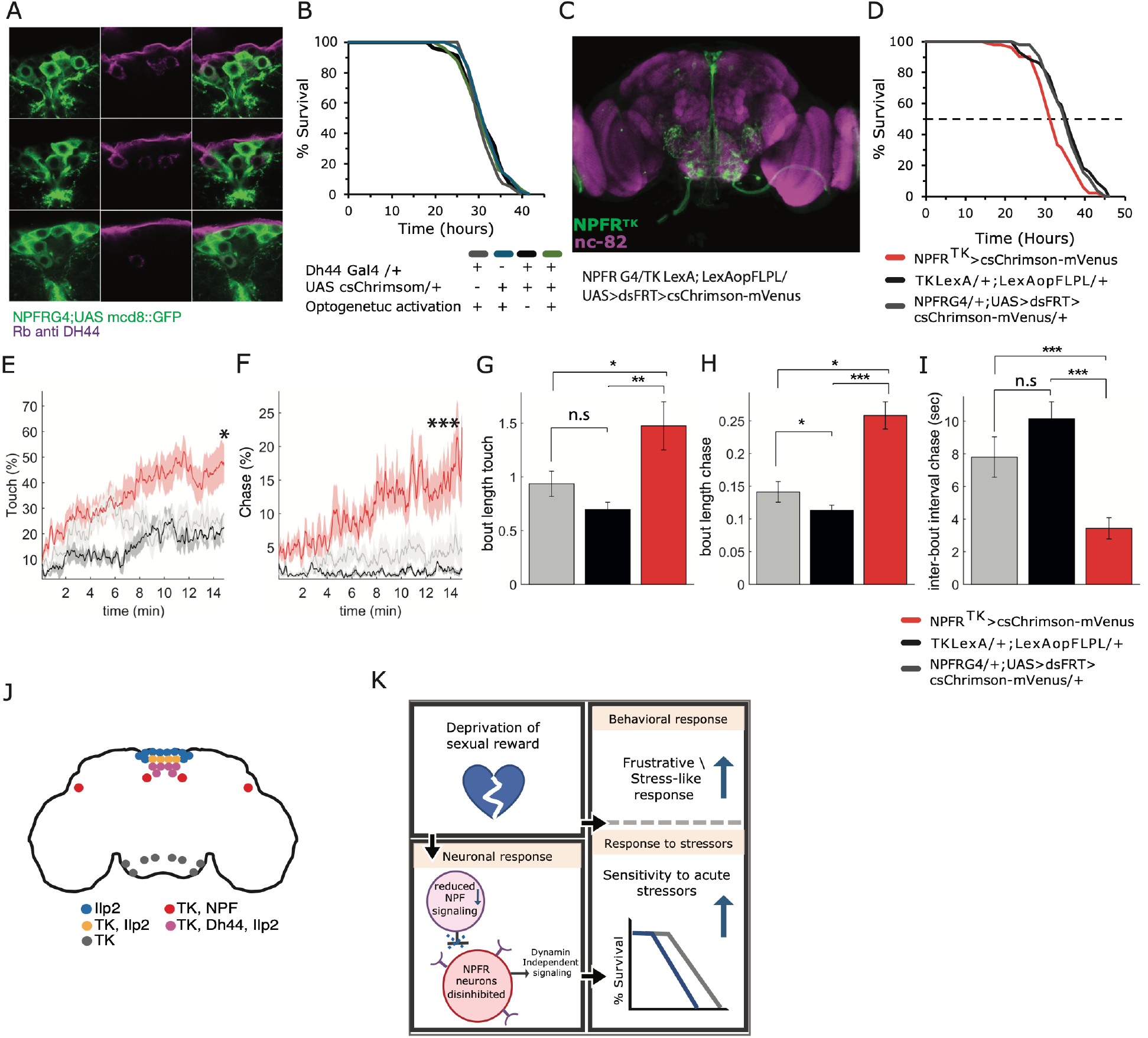
A subset of 12-16 NPFR neurons regulate both sensitivity to starvation and male-male social interaction. **A.** A portion of NPFR neurons co-express the neuropeptide DH44. Colocalization of DH44 using rabbit anti DH44 (magenta) in GFP expressing NPFR neurons. **B.** No effect for activation of Dh44 neurons on starvation resistance. *p*>0.05, n=56 for all cohorts. **D.** Shared NPFR^TK^ neurons as visualized using genetic interaction between NPFR and TK drivers: NPFRG4;+, TK-LexA;+, +;LexAop-FlpL, +;UAS<dsFRT>cs-Chrimson-mVenus flies. Green marks NPFR^TK^ neurons, magenta marks nc-82. **D.** Activation of NPFR^TK^ neurons enhances sensitivity to starvation (NPFR^TK^, red, n=51) compared to genetic control flies (TK-LexA;LexAop-FLPL, black, n=44; and NPFR G4;UAS<dsFRT>csChrimson-mVenus, gray, n=51). \**p*<0.05, \*\**p*<0.01. **E-I.** Analysis of male-male social interaction during optogenetic activation of NPFR^TK^ neurons compared to genetic controls using the FlyBowl system. Activation of NPFR^TK^ neurons induces higher rates of touch behavior (E), chase events (F), longer duration of touch events (G), longer duration of chase events (H), and higher frequency of chase as reflected by linger inter-bout intervals between chase events (I). n=13 for NPFR^TK^(red). n=13 TK-LexA;LexAop-FLPL (black), and n=12 NPFRG4;UAS<dsFRT>csChrimson-mVenus (gray). \**p*<0.05,\*\**p*<0.01, \*\*\**p*<0.001. ANOVA or Kruskal-Wallis with post-hoc Tukey’s or Dunn’s test, and FDR correction for multiple comparisons were performed. **J.** Schematic representation of brain NPFR neurons that intersect with TK, Ilp2, DH44, FoxO and NPF neurons. **K.** Summary of main findings according to which deprivation of sexual reward induces a stress-like response and increases sensitivity to subsequent acute stressors. This is mediated by a neuronal response: Sexual deprivation decreases NPF signaling, thereby disinhibits NPFR neurons and induces a dynamin-independent activity, which increases sensitivity to starvation.

Lastly, activation of a small neuronal population consisting of 22–26 cells that co- express the neuropeptide Tachykinin (NPFR^TK^, Fig. 5C), induced starvation sensitivity that was similar in its extent to the activation of the entire NPFR population (Fig. 5D). The effects do not depend on the TK neuropeptide itself, as its knockdown did not affect starvation rates (Supp. Fig. 7A, B), suggesting that activation by itself is responsible for the enhanced sensitivity. Remarkably, acute activation of these neurons mimics also the heightened arousal and some of the aggression-related features associated with the frustrative-like state. This included high ratios of male-male touch and chase events (Fig. 5E, F), higher frequency of chase events (Fig. 5G), higher persistence reflected by longer duration of touch and chase actions (Fig. 5H, I) and as a result, reduced formation of social clusters (Supp. Fig.8). In addition, the activation of NPFR^TK^ neurons resulted in increased coordination between pairs of flies as seen by lower values of features that measure relative changes in angle and speed between two close individuals (absanglefrom1to2, absphidiff and absthetadiff; Supp Fig. 7C), presumably suggesting that flies engage more persistently with others when interacting. In many cases the high arousal and persistence resulted in the formation of long chains containing 4–8 flies (Supp. Fig. 7C). Interestingly, we noticed that part of the NPFR^TK^ neuronal population co-expresses Dh44 and NPF (Supp. Fig. 7D, E), the activation of which does not contribute to the observed effects, therefore limiting the neurons regulating sensitivity to starvation and social responses to a subset of only 12–16 NPFR^TK^ neurons. Taken together, these results suggest that the disinhibition of a small subset of NPFR neurons in rejected males promotes behavioral responses that may serve to increase the odds of obtaining a sexual reward, and at the same time impair the ability of male flies to resist starvation stress.

## Discussion

In this study, we discovered that the NPF/NPFR system serves as a junction that integrates the crosstalk between reward and stress, and showed for the first time that *Drosophila* males perceive failures to obtain sexual reward as social stress. We used a collection of behavioral paradigms to explore responses altered by sexual interaction and discovered that repeated events of failure to mate lead to complex behavioral and physiological responses, encoded in part by the disinhibition of a subset of NPFR neurons. This includes avoiding interaction with other male flies and, at the same time, competing over mating partners via increased aggression and prolonged copulation (known as mate guarding); the latter is strengthened by the increased production of certain seminal fluid proteins that facilitate stronger post-mating responses in female flies. The regulation of ejaculate composition and mating duration were previously described in *Drosophila* males in response to perceived competition with rival male flies as a strategy to transfer a higher amount of accessory gland proteins (Acps) to intensify the females’ post-mating responses^47, 108, 109^. Although rejected males were not exposed directly to other male flies, the observed extension of copulation events and increased expression of *sex-peptide* suggest that failure to mate induces responses like those following the perception of competing male flies, presumably by evaluating the quality of their sexual interaction with female flies. It remains to be tested whether the enhanced investment in mating-related elements promotes mating success.

While most studies focus on mechanisms that encode reward on a scale of zero to one, our findings demonstrate the existence of negative values, where failure to obtain rewards is different than its lack, as shown by the clear differences between rejected and virgin males who never experienced mating or rejection. This provides a conceptual framework for investigating mechanisms that regulate deprivation or omission of an expected reward which is located at the negative part of the scale, when organisms fail to obtain a reward they expect to receive despite signals for its presentation^92^. This condition is known to induce a frustrative-like state, characterized by increased motivation to obtain the reward^93, 95, 99^, increased levels of arousal^99, 100^, agitation^94^, grooming^92, 100^, stress-associated behaviors^93, 94^, drug consumption^95, 101^, locomotion^94^, and aggression^95^. The use of courtship suppression to deprive male flies of the inherent expectation of a sexual reward provides a possible model for frustration-like stress responses in *Drosophila*. While courtship suppression is associated with reduced courtship and presumably a defeat-like state, we discovered that rejected males are rather persistent in their attempts to obtain a sexual reward. This finding is not completely surprising considering the innate nature of mating motivation and the presence of female aphrodisiac pheromones. It can also be attributed to measuring various courtship actions rather than to the overall percentage of time spent courting, which is the usual indicator for the quantification of courtship. In response to repeated failures to mate, rejected males exhibit features characteristic of a frustration-like state, such as persistent mating actions (some of which are elongated), increased arousal, increased aggression, as well as longer mating duration upon successful mating encounters, all of which are reminiscent of a high motivational state.

While this motivational state may assist males in coping with social challenges associated with failure to mate, it is perceived as a stressful experience manifested by temporal costs in the form of sensitivity to acute stressors. This is the first example in flies that not obtaining a sexual reward is not simply a case of reward deprivation but also a stressful experience. But what mechanism governs the behavioral and physiological responses to deprivation of sexual reward? In search of mechanisms that could explain the behavioral and physiological responses to the failure of obtaining a sexual reward, we examined three possible directions: (1) sleep and activity (2) energetic costs or modulation of metabolic pathways, and (3) disinhibition of NPF target neurons.

Regarding the possibility the high motivational state is associated with high energetic costs, and although Gendron et al.^75^ showed that exposure to female pheromones lowers body triglyceride levels in males, we did not detect metabolic changes that could explain sensitivity to starvation or oxidative stress. Still, our findings that some metabolites, such as 5-Aminolevulinic acid, are enriched in rejected flies may indicate a reduction in Heme synthesis and, consequently, an elevation in protoporphyrin, as well as a possible reduction in heme oxygenase (HO). While there is some evidence that protoporphyrin can act as an antioxidant^110, 111^, it mainly functions as a pro-oxidative agent^112, 113^. HO is a rate-limiting enzyme that degrades heme into biliverdin, carbon monoxide (CO), and iron^114^. In *Drosophila*, the *ho* gene is expressed in different brain tissues, including in the optic lobe, central brain, and glial cells, and plays an important role in cell survival and protection against paraquat-induced oxidative stress^115^. This is consistent with rejected males experiencing increased sensitivity to oxidative stress, reflected in their heightened sensitivity to paraquat. However, this does not explain why rejected males were also more sensitive to starvation. Five of the metabolites were specifically enriched or depleted in rejected compared to both mated and naïve-single cohorts: glycine, tryptophan, 5-aminolevulinic acid (5ALA), acetyl-glutamine, and stearic acid (Supp Fig. 5E, Table S3). Though tryptophan can be converted to serotonin or tryptamine, and most of the tryptophan metabolizes to kynurenine pathway (KP) metabolites, which contribute to a shorter lifespan^116–119^, we did not document any significant accumulation or depletion of metabolites in the KP, or a difference in the abundance of serotonin (Supp Fig. 5E). Similarly, no difference was observed in oxidative agents or in antioxidant activity and glucose levels, though trehalose (the main circulating sugar in flies) levels declined in naïve-single males compared to mated and rejected males, indicating a possible change in insulin signaling^120–123^.

After eliminating the contribution of sleep or metabolic deficit to the inability of rejected males to cope with stress, we demonstrated that their sensitivity to starvation stress is caused by the disinhibition of NPF target neurons, mimicked in our experiments by optogenetic activation of NPFR neurons as well as knock-down (KD) of *npfr*. We mapped the relevant neurons to a subset of 22-26 NPFR^TK^ neurons, the activation of which is sufficient to induce sensitivity to starvation and some of the behavioral phenotypes observed in rejected males. The identified NPFR sub-population can be divided into smaller known subpopulations: six of these neurons express the CRF-like Dh44, and two pairs of NPFR^TK^ neurons, L1-l and P1, colocalize with NPF-expressing neurons. While CRF neurons in mammals mediate reward-seeking behaviors and responses to social stress^13, 14, 124–130^ and the fly Dh44 neurons are known to facilitate aggressive behaviors^131^, we did not find evidence to support their role in regulating responses to social stress. We concluded that the activation of a subset of 12-16 neurons co-expressing the Tachykinin neuropeptide governs both starvation sensitivity and enhanced arousal and some features of aggressive behaviors. It should be noted that the effects of optogenetic activation of NPFR^TK^ neurons was tested using the FlyBowl setup, a context that does not allow for the expression of lunging behavior (due to low ceiling) but do support the expression of other forms of aggression such as chases and wing threats. Therefore, the activity of the NPF\NPFR circuit alone is sufficient to facilitate a stress-like response caused by the deprivation of a reward and increase subsequent sensitivity to acute stressors.

While the subset of NPFR neurons that mediate the behavioral and physiological response shares expression with the *Tachykinin*^61^, the TK neuropeptide itself does not play a role in modulating the sensitivity to starvation upon sexual rejection. In addition, while Wohl et al., ^132^ found that TK supports cholinergic synaptic transmission, our results suggest that the increased sensitivity to starvation is mediated by a dynamin-independent signaling mechanism, possibly bulk endocytosis for vesicle retrieval or neuropeptide release, which commonly does not involve synaptic vesicle reuptake. Further dissection of the NPFR and NPFR^TK^ neuronal subpopulations is needed to better understand the role of each cell type in response to sexual reward deprivation.

This is the first documentation of a frustration-like stress response in *Drosophila*, which is caused by the innate expectation of a natural reward that is not achieved. Our findings offer a paradigm shift in the perception that rejected males refrain from obtaining a sexual reward by showing that their motivation to copulate increases, not declines, and thus can serve as a conceptual framework to study the interplay between reward-seeking behaviors such as ethanol consumption, stress, and addiction. The bi-directional role of the NPF system in mediating reward and suppressing response to stressors^59^ and the remarkable similarity to mammalian NPY^133^ suggest that it is possible to simplify the complexity of studying the crosstalk between reward, stress, and reproduction in mammals by stripping it down to its most fundamental building blocks in fruit flies.

## Materials and Methods

### Fly lines and culture

*Drosophila melanogaster* WT Canton S flies were kept at 25C°, ∼50% humidity, light/dark of 12:12 hours, and maintained on cornmeal, yeast, molasses, and agar medium. Most fly lines were backcrossed to a Canton S background. UAS UAS-*unc84*-2XGFP and UAS mCD8 GFP were obtained from HHMI Janelia Research Campus. For INTACT, UAS-*unc84*-2XGFP transgenic flies were crossed with NPFR-GAL4 flies. NPFR-Gal4, UAS-NPFR RNAi, NPF-GAL4 flies were a gift from the Truman lab (HHMI Janelia Campus). UAS<dsFRT>cs-Chrimson-mVenus in attp2, and LexAop-FLPL flies were a gift from the Heberlein lab (HHMI Janelia Campus). The following lines were ordered from the indicated fly centers: TK-LexA (Bloomington #54080), UAS-tk-HA (ORF F000997), UAS-tk RNAi (VDRC #103662), Dh44-Gal4 (VDRC #207474).

### Sexual experience protocols

Males and females were collected within 2 h of eclosion on CO2, 3-4 days before courtship conditioning. Males were collected into narrow glass vials (VWR culture glass tubes 10X75mm) containing food and kept single housed until the conditioning. To generate mated females for the experiment, mature males were added to the females ∼16 h before the experiment. All flies were kept in the incubator at 25°C, ∼50% humidity, and light/dark of 12:12 hours. The mated females were separated from the males on the morning of the conditioning. During the conditioning, the temperature was kept at about 25°C, and humidity was ∼55%.

#### Generation of rejected males

Individual males were placed with mated females for 3 one-h conditioning trials (separated by 1-h rests) a day for two or four consecutive days. Females were removed after each trial. Males from the rejected cohort that managed to mate and males from the mated cohort that did not end up mating during all sessions were discarded. At the end of each session, the female fly was removed, and the males that experienced rejection were kept in the original vial for one hour of rest. Males were monitored every 10 minutes to ascertain lack of mating, and when mentioned the number of males exhibiting courtship action during training sessions was documented.

#### Generating mated males

To generate the “mated-grouped” cohort, individual males were housed with virgin females for 3 one-h conditioning trials (separated by 1-h rests) a day for two or four consecutive days. Females were removed after each trial.

#### Generating Naïve-single males

irgin males were collected within 2 h of eclosion and kept separately in small food vials during the entire trial. Gentle handling was performed parallel to rejected and mated males conditioning sessions. The naïve-single male cohort was kept in the behavior chamber during the training phase, and the vials containing the males were handled as similarly as possible to the rejected and mated cohorts, without inserting female trainers. Detailed protocol is previously described^86^.

### Behavioral, molecular, and metabolic analysis following experience phase

#### Analysis of social group interaction using the FlyBowl system

At the end of four days of sexual experience, rejected, mated and naive-single male flies were inserted in groups of 10 into Fly Bowl arenas^89^, and their behavior was recorded for 30 minutes and analyzed using CTRAX, FixTrax^88^ and JAABA^89^. For kinetic features, scripts were written in MATLAB to use the JAABA code to generate the statistical features as specified in Kabra et al,^89^. Time series graphs (per frame) were created using JAABA Plot^89^. Quantification of complex behaviors was done using JAABA Classifiers^89^ to identify specific behaviors: Walk, Stop, Turn, Approach, Touch, Chase, Chain, Song, Social Clustering, and Grooming. Bar graphs were created using JAABA Plot^89^. Network analysis was performed using an interaction matrix according to the interaction parameters described previously^88^. Two interaction matrices were created for each movie, one with the total number of frames each pair of flies were interacting divided by the number of frames in the movie and another with the number of separate interactions between each pair of flies divided by the maximum number of possible interactions, calculated as:

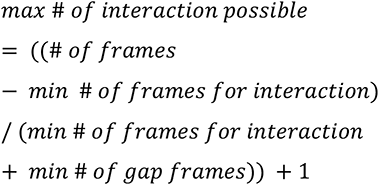

The parameters to define an interaction are angle subtended by the other fly > 0, distance between the nose of the current fly to any point on the other fly ≤ 8 mm, number of frames for interaction ≥ 60 and number of gap frames ≥ 120. Interaction end is defined when distance or angle conditions are not maintained for 4 seconds. Networks and their features were generated from the interaction matrix in R using the igraph package. The function that was used to generate the networks is “graph_from_adjacency_matrix” with parameters “mode = undirected” and “weighted = TRUE”. Density was calculated on all movies with the formula:

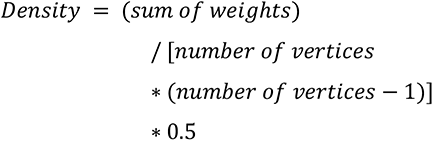

Modularity was calculated using the “modularity” function on output from the “cluster_walktrap” function65. Strength was calculated using “strength” function and SD Strength was calculated on all movies using “sd” function on the strength value. Betweenness Centrality was calculated on all flies using the “betweenness” function and SD Betweenness Centrality was calculated on all movies using “sd” function on the Betweenness Centrality value. Box plots were created using R. Each feature of the FlyBwol experiment was standardized according to all values calculated in our experiments for that feature to generate a z-score. Scatter plots were created using R.

#### Aggression assay

At the end of four days of sexual experience, pairs of rejected or mated male flies were introduced into aggression arenas (circular chambers, about 0.08 cm3 in volume), which contained a mixture of agarose and apple juice (1% agarose, 50% apple juice) that was placed in arenas to enhance aggressive behavior. Flies were filmed for 30 min with Point-Grey Flea3 (1080×720 pixels) at 60 fps. Aggressive behavior was later quantified by counting the number of lunges for each pair using CADABRA software^134^. The log2 ratio between the number of lunges in rejected and mated flies was calculated for each pair, and then a one-sample t-test was performed to test whether the mean ratio is significantly different from 0.

#### Copulation duration

At the end of four days of sexual experience, Rejected and naive-single male flies were inserted into courtship arenas (circular chambers, about 0.04 cm3 in volume) with virgin females and were allowed to mate for 1 hour. They were recorded for the whole experiment using a Point-Grey firefly camera. Courtship arenas consist of 25 flat arenas each arena containing only one pair of male-female flies. The copulation duration was measured from the moment the mating began until it ended. We calculated the time in seconds for each fly and the average for each group.

#### Quantitative Real-Time PCR analysis

At the end of 4 days of sexual experience male flies were flash frozen. Frozen flies were placed on ice and decapitated using a scalpel. Total RNA was extracted from ∼15 frozen bodies, using TRIZOL reagent according to the manufacturer’s protocol. mRNA was reverse transcribed using the BIORAD cDNA synthesis kit. cDNA was analyzed by quantitative real-time PCR (BIORAD CFX96) using specific primers for the head and for the body. Relative expression was quantified by ΔΔCT method using RPL32 as a loading control. We run each sample in triplicates. Each experiment was repeated four times using independent sets of experimental flies.

#### Sensitivity to starvation stress

Following two days of sexual experience or when mentioned after optogenetic activation, males were transferred to glass vials containing 1% agarose and were kept singly throughout the experiments. The number of live flies were recorded every couple of hours.

#### Sensitivity to oxidative stress

Following two days of sexual experience, male flies were introduced singly within glass vials containing standard food supplemented with 20mM paraquat (856177, Sigma-Aldrich), and their survival curve was monitored every 2 hours.

#### Longevity

Following two days of sexual experience, males were transferred to vials containing food. Males were kept in isolation and the number of living flies was recorded every day. Flies were transferred to new vials twice a week. Log-rank or Renyi-type test (REF) with FDR correction was performed.

#### Sleep\locomotion assays

Following two days of experience mated, rejected and naïve-single male were placed in a 48-well cell and tissue culture plate (TC plate) by gentle aspiration. Each well contained 1% agar, 7% sucrose, and 0.7% yeast extraction. Locomotor data were collected using DanioVision (Noduls) software and raw data files were analyzed with EthoVision XT software. Activity was measured in 1-min bins and sleep was identified as 5 min of consolidated inactivity, defined as no movement^135, 136^. Sleep and activity data were analyzed using MATLAB. The training was copleted at ZT5.5 and sleep was assessed from ZT6 onwards in the DanioVision (Noduls) system owing to the time spent in introducing flies into individual wells.

#### TAG, Glucose levels evaluation

TAG levels were assessed as described^75^ with modifications: After two days of sexual experience, experimental males were divided into groups of 5 and were homogenized together in 100 μl NP40 substitute assay reagent from Triglyceride colorimetric assay kit 10010303 (Cayman JM-K622-100). Homogenate was centrifuged at 10,000 x g for 10 min at 4°C, and the supernatant was collected. Triglyceride enzyme mixture (10010511) was used to hydrolyze the triglycerides and subsequently measure glycerol by a coupled enzymatic reaction. TAG concentrations were determined by the absorbance at 540nm and estimated by a known triglyceride standard. The absorbance was measured using a SynergyH1 Hybrid Multi-Mode microplate Reader. Body and hemolymph glucose were extracted as described^137^. Briefly (with modifications):

Whole bodies: After two days of sexual experience, 5 males were placed in each sample tube and weighed using Fisher scientific ALF104 analytical balance scale. Then, flies were homogenized in 100 ml cold PBS on ice. The supernatant was heated for 10 min at 70°C, then centrifuged for 3 min at maximum speed at 4°C. The supernatant was collected and transferred to a new 1.5 ml tube. Hemolymph: After courtship conditioning, males were sedated on ice and carefully punctured in the thorax using sharp stainless-steel tweezers. 40 punctured flies were placed in each 0.5 ml microfuge tube with a hole at the bottom made by a 25G needle. The 0.5 microfuge tube was then placed in a 1.5 ml tube and centrifuged at 5000 rpm for 10 min at 4°C. Hemolymph was collected, and samples were heated for 5 min at 70°C. Glucose was measured using High sensitivity Glucose Assay kit (MAK181 Sigma-Aldrich). Glucose concentration is determined by coupled enzyme assay, which results in fluorometric (lex = 535/lem = 587nm) products and was assessed using SynergyH1.

#### Metabolite extraction and LC-MS metabolomic analysis

After two days of sexual experience, males were flash-frozen and decapitated using a microscalpel. For each sample, 5 heads were transferred into soft tissue homogenizing CK 14 tubes containing 1.4 mm ceramic beads (Bertin corp.) prefilled with 600 ul of cold (−20 °C) metabolite extraction solvent containing internal standards (Methanol: Acetonitrile: H2O:50:30:20) and kept on ice. Samples were homogenized using Precellys 24 tissue homogenizer (Bertin Technologies) cooled to 4 °C (3 × 30 s at 6000 rpm, with a 30 s gap between each cycle). Homogenized extracts were centrifuged in the Precellys tubes at 18,000 g for 10 min at 4 °C. The supernatants were transferred to glass HPLC vials and kept at −75 °C prior to LC-MS analysis. LC-MS analysis was conducted as described^138^. Briefly, Dionex Ultimate ultra-high-performance liquid chromatography (UPLC) system coupled to Orbitrap Q-Exactive Mass Spectrometer (Thermo Fisher Scientific) was used. The resolution was set to 35,000 at a 200 mass/charge ratio (m/z) with electrospray ionization and polarity switching mode to enable both positive and negative ions across a mass range of 67–1000 m/z. UPLC setup consisted of ZIC-pHILIC column (SeQuant; 150 mm × 2.1 mm, 5 μm; Merck). 5 µl of cells extracts were injected and the compounds were separated using a mobile phase gradient of 15 min, starting at 20% aqueous (20 mM ammonium carbonate adjusted to pH 9.2 with 0.1% of 25% ammonium hydroxide):80% organic (acetonitrile) and terminated with 20% acetonitrile. Flow rate and column temperature were maintained at 0.2 ml/min and 45 °C, respectively, for a total run time of 27 min. All metabolites were detected using mass accuracy below 5 ppm. Thermo Xcalibur 4.1 was used for data acquisition. The peak areas of different metabolites were determined using Thermo TraceFinder^TM^ 4.1 software, where metabolites were identified by the exact mass of the singly charged ion and by known retention time, using an in-house MS library built by running commercial standards for all detected metabolites. Each identified metabolite intensity was normalized to ug protein. Metabolite-Auto Plotter^139^ was used for data visualization during data processing.

#### Detailed analysis of courtship suppression

WT males and females were collected on CO2 3-4 days before the recording. Males were kept in groups of 25 per vial. To generate mated females for the experiment, virgin females were introduced to males ∼16 hours before the experiment. All flies were kept in the incubator at 25°C, ∼50% humidity, and light/dark of 12:12 hours. Prior to the conditioning, the mated females were separated from the males on CO2 on the morning of the recording. During the recording, the temperature was kept at ∼25°C, and humidity was ∼55%. Since the extent of courtship display is shaped by circadian rhythmicity, where male flies depict the highest courtship activity closest to the onset of light, and their general activity declines towards noon, the first session started right after the onset of light, and the other two sessions took place in the afternoon. Virgin male flies were exposed to either mated or virgin female for three one-hour sessions, and their behavior was recorded using a Point Grey Firefly camera and analyzed in detail during the first 10 minutes of each interaction. At the end of each session, female flies were removed, and the males that experienced rejection were kept isolated in narrow glass vials for one hour. At the end of the rest hour, males were returned to their original location in the courtship arena for the recording. To compare the courtship behavior of rejected and control males, virgin males from the control cohort were replaced at the beginning of each session. Different aspects of courtship behavior were analyzed manually using “Lifesong” software. Courtship index for a given male is the fraction of time a male fly spent in courtship activity in the 10 min observation period (600 sec). It is calculated by dividing the number of seconds the male courted over the total observation time (CI = courtship behavior [sec] · 100 / total observation [sec]).

#### Optogenetic Activation of NPFR, Dh44, and NPFR^TK^ neurons

Light-induced activation of red-shifted Channel Rhodopsin UAS-CsCrimson was achieved by placing glass fly vials containing one fly each over red LEDs (40 Hz, 650nm, 0.6 lm @20mA). Activation protocol consisted of 3x5 min-long activation periods spaced by 1 h and 55 min resting intervals for 2 consecutive days.

#### Neuronal activation combined with inhibition of synaptic vesicle release

Flies expressing Cs-Chrimson and UAS-Shibire^ts^ in NPFR neurons were subjected to one of four conditions for two days: (1) Three 5-min-long optogenetic activations spaced by 1 h and 55 min resting intervals (under constant dark) at constant 18-20°C served as a positive control. (2) Three 10-min-long sessions at 28-29°C under constant dark followed by 5 min-long optogenetic activations spaced by 1 h and 45 min resting intervals at 18-20°C, also under constant dark. (3) Three 15-min-long sessions at 28-29°C under constant dark, spaced by 1 h and 45 min at 18-20°C, also under constant dark, served as synaptic release block control. (4) Flies kept at a constant 18-20°C and constant dark served as a negative control. After the last activation, flies were transferred into glass vials containing 1% agarose.

#### Optogenetic activation using the FlyBowl setup

Flies expressing CsChrimson in various neuros were introduced into the FlyBowl arenas and their behavior under optogenetic illumination was recorded as described in Bentzur et. al^88^. In brief: groups of 10 male flies, which were socially raised in groups of 10 for 3-4 days, were placed in FlyBowl arenas, and their behavior was recorded at 30 fps for 15 min and tracked using Ctrax^90^. Automatic behavior classifiers and Per-frame features were computed by JABBA^89^ tracking system. Data of all behavioral features were normalized to the percentage of difference from the average of each experiment for visualization. Details about the different features are found in Figure S4.

#### Immunostaining

Whole-mount brains were fixed for 20 min in 4% paraformaldehyde (PFA) or overnight in 1.7% PFA. Preparations were blocked for 1h at 4°C with gentle agitation in 0.5% BSA, and 0.3% Triton in PBS. The following primary antibodies were used: Rabbit anti-GFP (LifeTech 1:500), the neuropile-specific antibody NC82, (1:50, The Jackson Laboratory), mouse anti-GFP (1:100, Roche), Rabbit anti-Dh44 (0.6:100), rabbit anti dILp2 (1:100, a kind gift from Takashi Nishimura lab), rabbit anti tk (1:1000 a Kind gift from Wei Song) were incubated overnight at 4°C. Secondary antibodies, goat anti mouse-Alexa488 (1:200-1:100), goat anti rabbit-Alexa568 (1:200-1:100), goat anti mouse-Alexa568 (1:1000) and goat anti rabbit-Alexa488 (1:1000) were incubated for 2hr at 4°C. DAPI (1:20). The stained samples were mounted with SlowFade^TM^ Gold antifade reagent (Thermo Fisher Scientific) and visualized using a Leica SP8 confocal microscope.

#### Statistical analysis

Data of each behavioral feature per experiment were tested for normality, and consequently, normally distributed data were tested by student’s t-test, one-way ANOVA followed by Tukey’s post-hoc. Non-parametric data were tested by Mann-Whitney or Kruskal-Wallis tests followed by Dunn’s or Friedman’s post-hoc tests. FDR correction for multiple comparisons was performed for all Flybowl experiments features. Statistical overrepresentation was generated using _PANTHER_140,141 (http://pantherdb.org/citePanther.jsp). Kmeans clustering method was performed (k= 3) to generate a heatmap of differentially expressed genes in NPFR neurons. Starvation resistance and longevity experiments were tested by Log-rank or Renyi-type test^142^ using R package version 3.2-11. FDR correction for multiple comparisons was performed for experiments with more than two experimental groups.

## Author Contributions and Notes

U.H, G..SO, D.R.N and J.R designed research, J.R, L.O, Y..KK, I.A, B.A and M.L performed research, L.B.B wrote software, J.R, E.G, I.A and H.P analyzed data; and J.R, U.H, D.R.N and G..SO wrote the paper. The authors declare no conflict of interest. This article contains supporting information online.

## Acknowledgments

We thank all members of the Shohat-Ophir lab for fruitful discussions and technical support. We would also like to thank Jennifer I. C. Benichou for statistical consultation. This work was supported by the Israel Science Foundation Grants 384/14 and 174/19.

---

**Figure S1.**
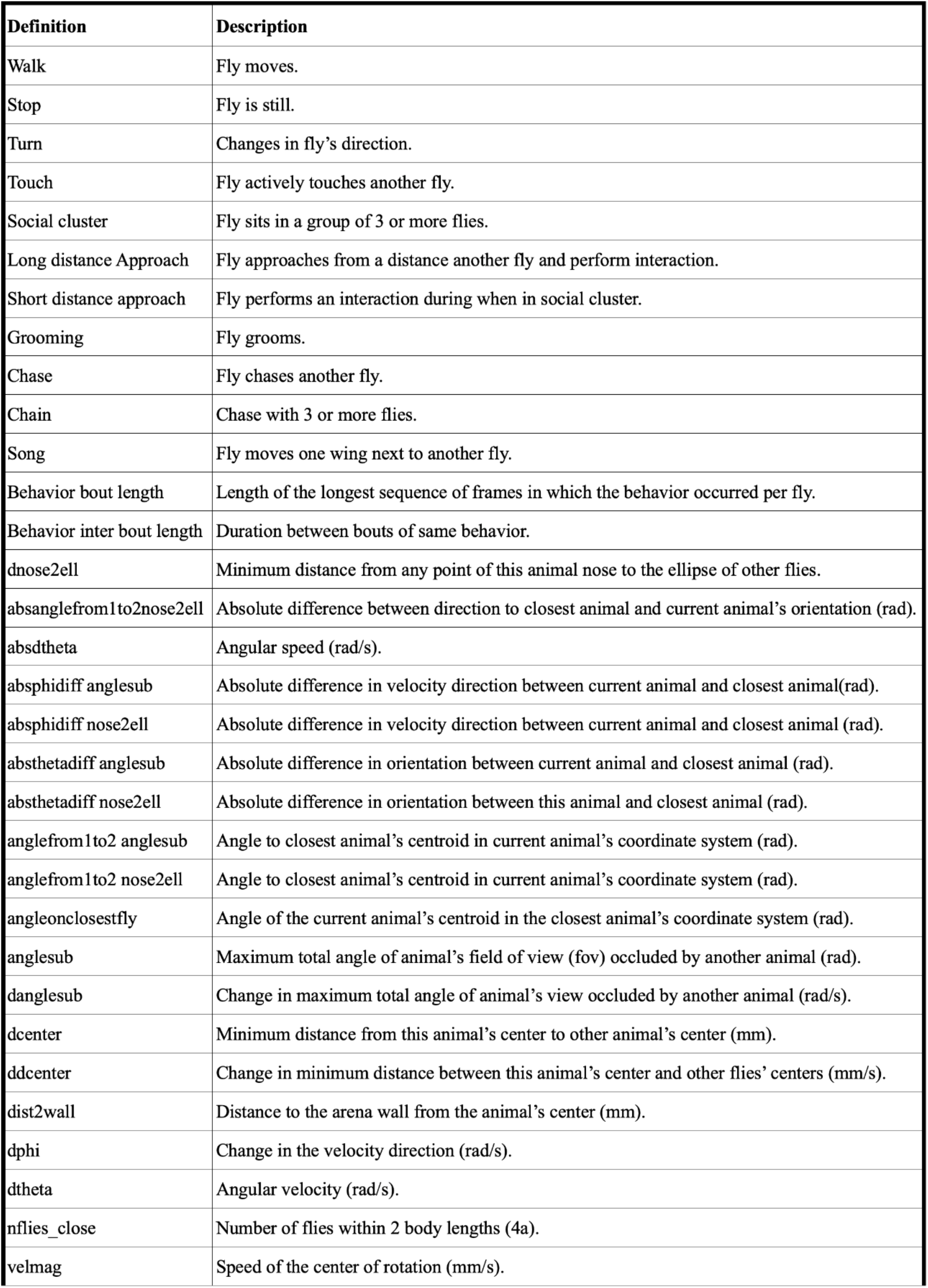
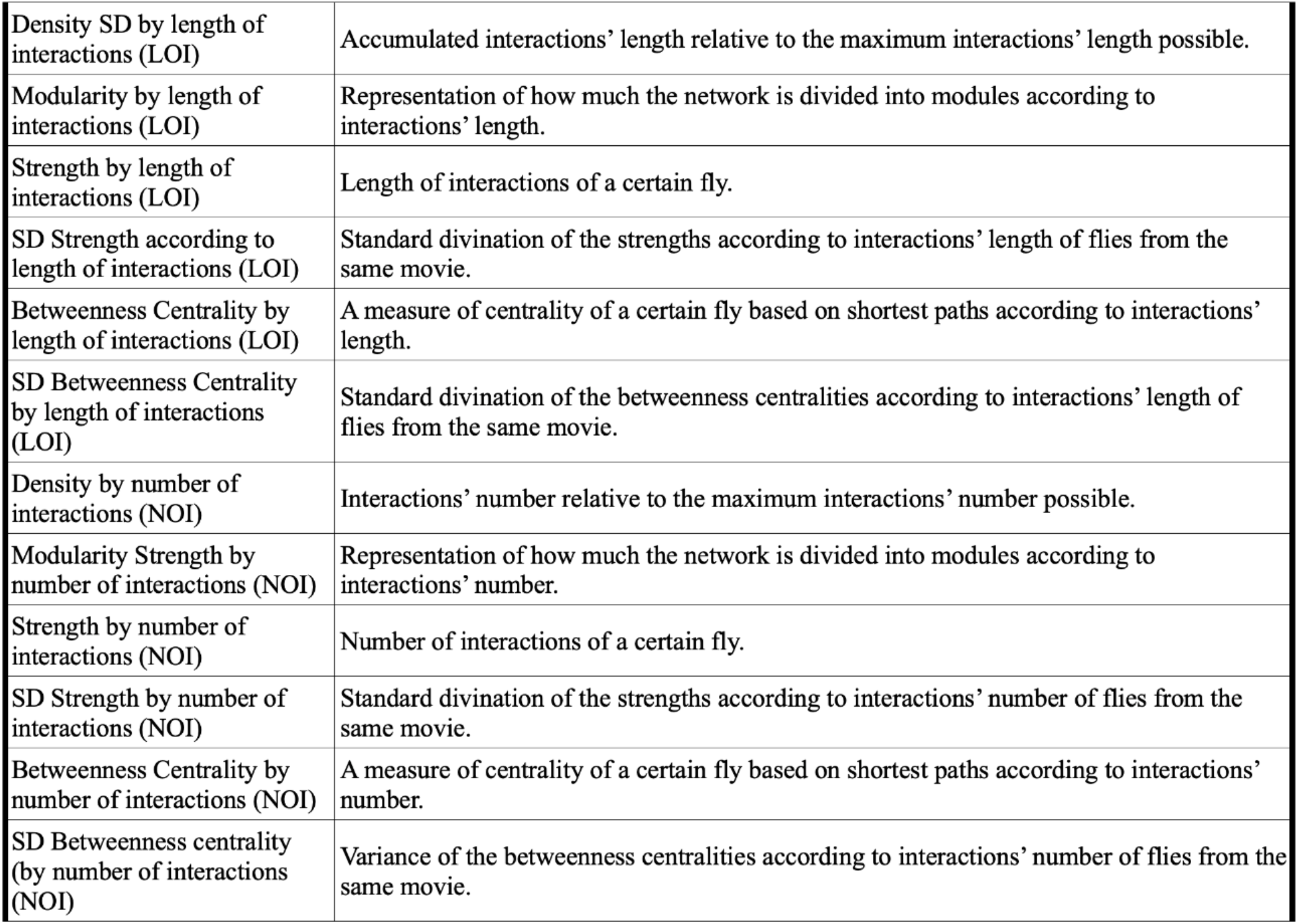
List of behavioral features presented in Figures 1, 2, 5, S2, S6 and S7.

**Figure S2.**
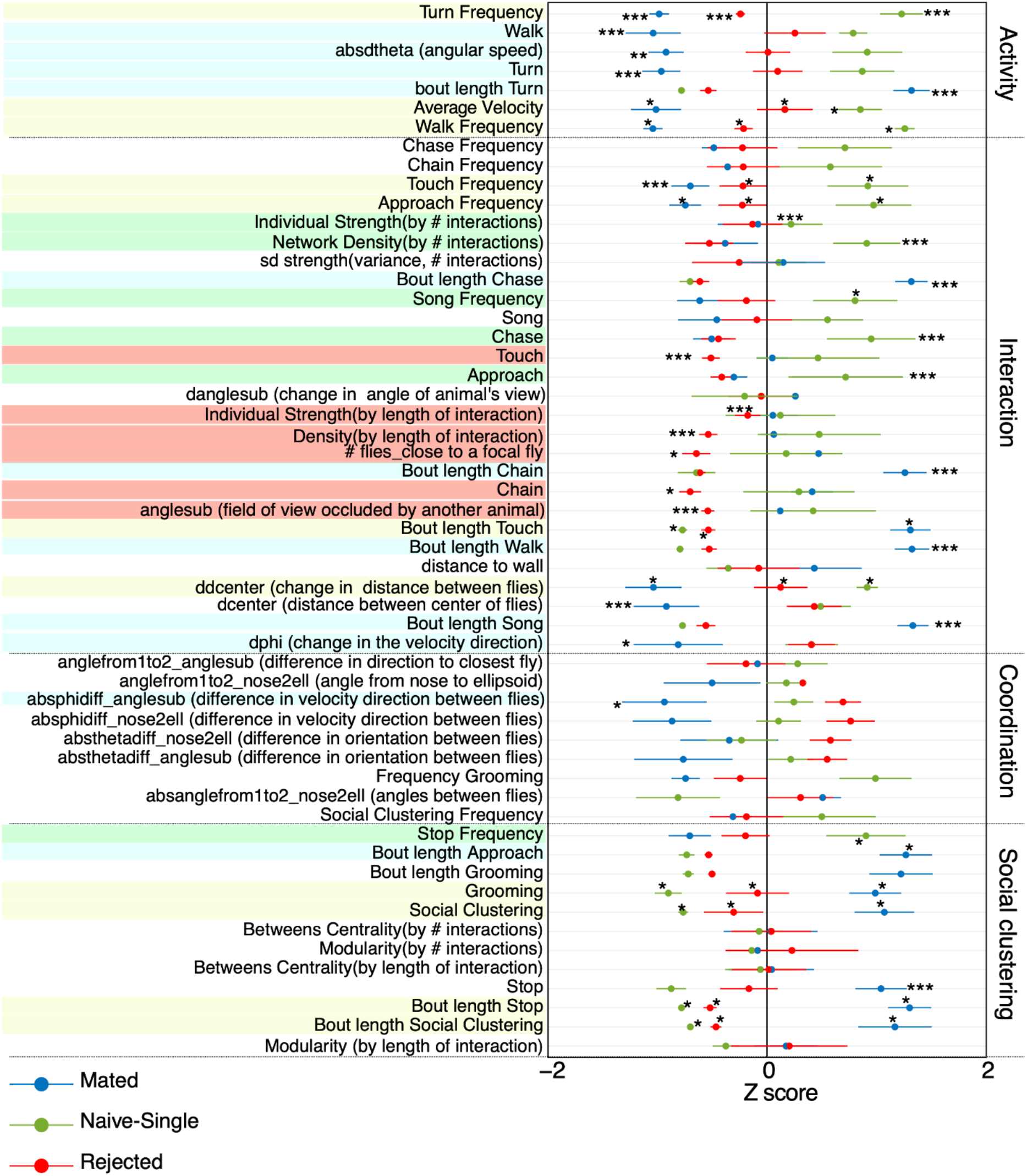
Behavioral signatures of mated, rejected, and naïve WT male flies. Data is represented as normalized Z scores of 60 behavioral parameters, n = 15, 10, 15 for mated, naïve-single, and rejected respectively. Statistical significance was determined by one-way ANOVA followed by Tukey’s range test for experiments that were distributed normally, and by Kruskal–Wallis test followed by Wilcoxon signed-rank test for experiments that were not distributed normally. FDR correction was used for multiple tests. LOI: calculated according to the length of interactions. NOI: calculated according to the number of interactions. Features marked in yellow exhibit statistically significant differences among the 3 cohorts. Features colored in blue are different between mated and the two other conditions. Features colored in red are different between rejected and the two other conditions. Features colored in green are different between naive and the two other conditions.

**Figure S3.**
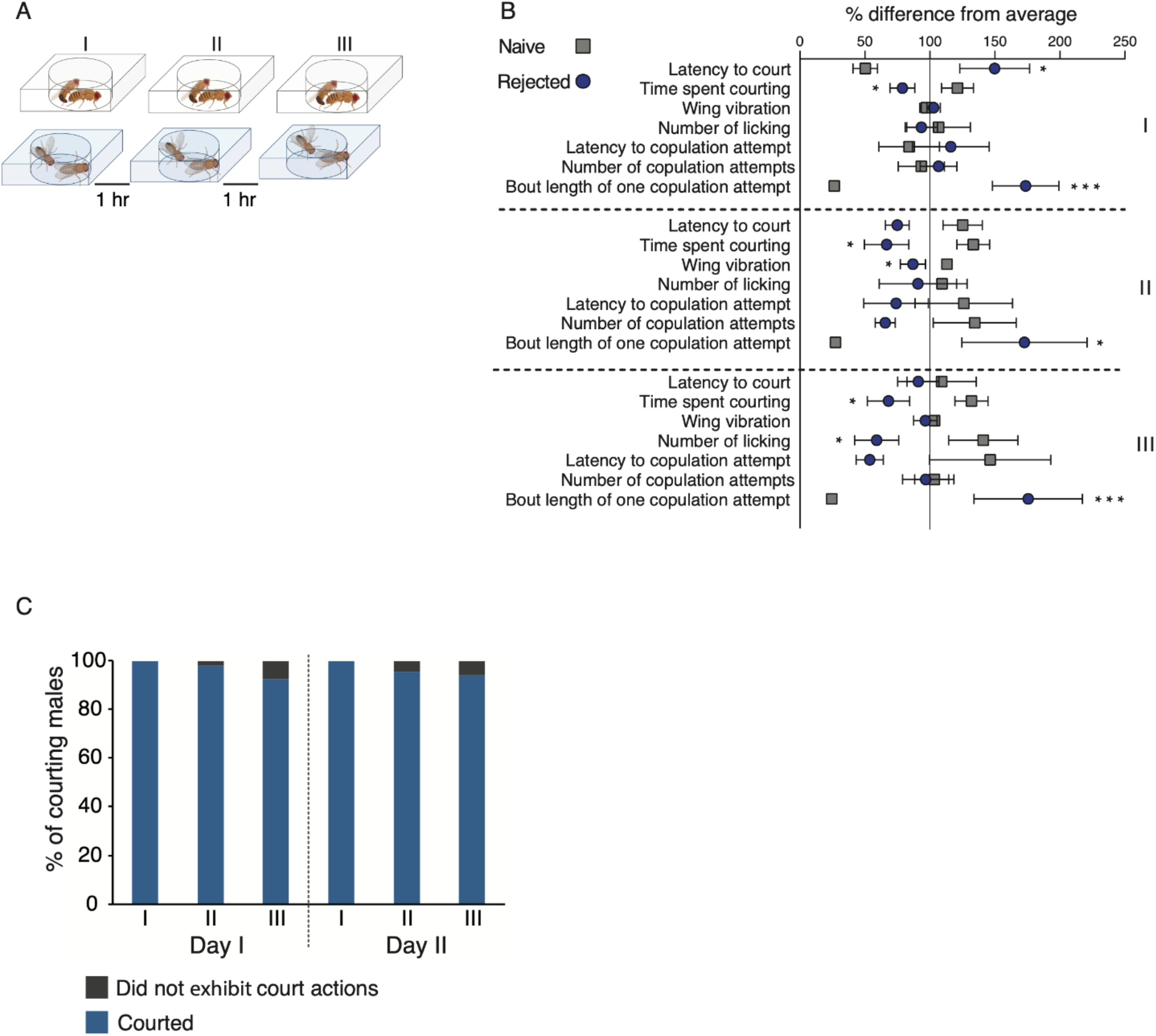
**A.** Schematic representation of courtship suppression analysis along repeated exposure to mated females. Virgin male flies were exposed to either mated or virgin females for three 1h sessions, and their behavior was recorded. At the end of each session, the females and males from the naive cohort (top) were removed, and the males that experienced rejection (bottom) were kept isolated in narrow glass vials for 1h. **B.** % difference from average courtship behaviors performed by rejected (blue circle) and naïve-single (gray square) males in the first (I), second (II), and third (III) sessions. Student’s t-test or Mann-Whitney was performed with FDR correction for multiple comparisons. \**p*<0.05, \*\*\**p*<0.001. **C**. Naive males were repeatedly introduced to sexually non-receptive females over two consecutive days. The number of courting males was documented for each session. Males that did not initiate courtship, or that succeeded to copulate were excluded from further analysis. Total n of males for day I= 97 first session (97 courted), 97 second session (95 courted), 95 third session (88 courted), n for day II= 86 first session (86 courted), 86 second session (82 courted), 82 third session (77 courted). Bar graph represents % of courting males in each session.

**Figure S4.**
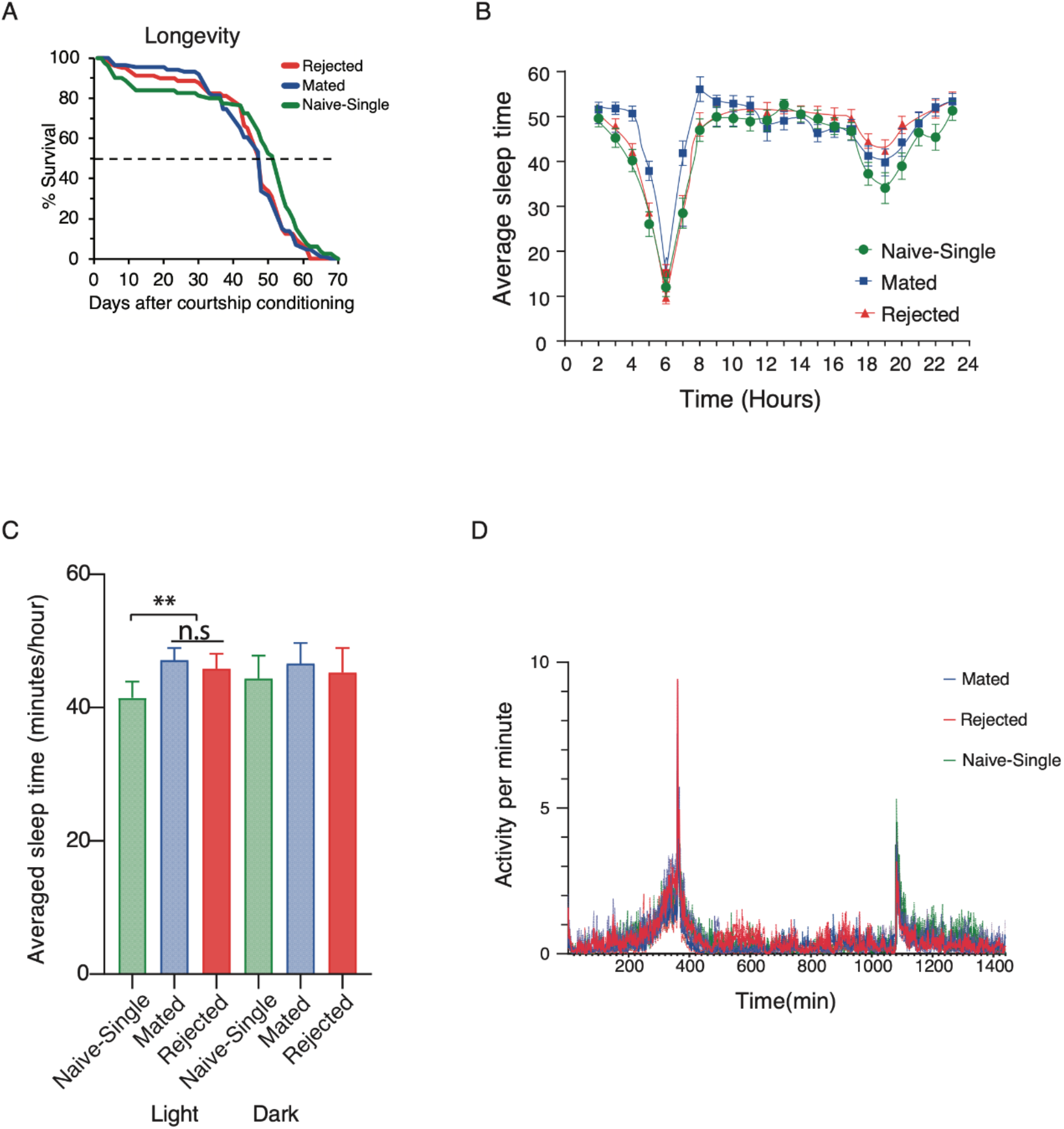
Rejected males did not exhibit a difference in sleep and activity patterns compared to both mated and naïve-single males. **A.** Longevity assay: single males (yellow, n=80) compared to mated (red, n=86) and rejected (blue, n=80) males. \**p*<0.05. Rényi test with PDF corrections for multiple comparisons was performed. **B-D**. Rejected males do not exhibit a difference in sleep and activity compared to both mated and single males. Sleep profiles depicting sleep amounts in 60 minutes binned intervals for single (green circles), mated (blue rectangles) and rejected (red triangles) males. Sleep and activity profile of naïve-single (green), mated (blue) and rejected (red) males. **B** and **C**. Sleep behavior illustrated as a sleep profile depicting sleep amounts in 1-hour binned intervals (B) and a stacked bar chart showing total sleep amount (C). Naive-single males exhibited decreased amounts of sleep during the light phase compared to mated and rejected males (p<0.01), no significant difference was observed between rejected and mated males (P>0.05). **D**. Activity data represented as 1-minute binned amount of movement. Mated males exhibited decreased activity compared to rejected and naive-single males (p<0.0001). No significant difference was observed between rejected and naive-single males (p>0.05). Repeated measures ANOVA with Tukey’s multiple comparisons test was performed in both A and B. n= 32(naïve-single, rejected), 31(mated).

**Figure S5.**
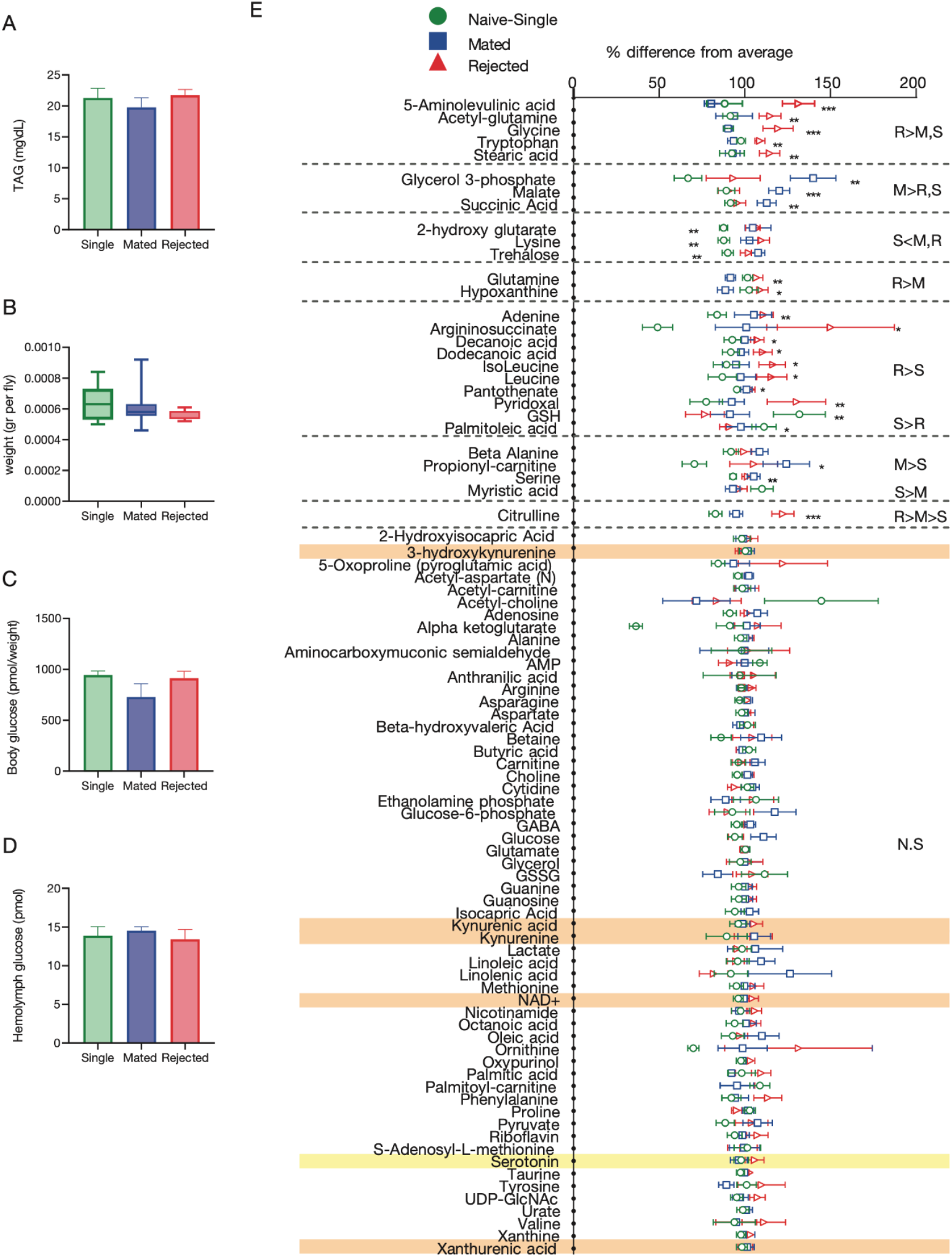
Courtship conditioning did not affect TAG and glucose levels and most head metabolites in males. **A-D.** Metabolic indices of rejected males (blue) compared to naïve-single (green) or mated (red) males. No differences were observed for measurements of (**A**) triglycerides (TAG, n=11 for all groups, 5 males/ sample). (**B**) weight (n= 10 naïve-single, 9 mated, 9 rejected, 5 males/sample). (**C)** hemolymph, or (**D)** body glucose (n=3 for all groups, 5 males/body sample, and 40 males/hemolymph sample). ANOVA or Kruskal-Wallis with post-hoc Tukey’s or Friedman test were performed. NS *p*>0.05. **E.** % difference from average (peak area/ total measurable ions) of metabolites detected using LC-MS in rejected, mated, and naïve-single males’ heads. 5-aminolevulinic acid, acetyl-glutamine, glycine, tryptophan, and stearic acid levels were higher in rejected males’ heads (blue triangles, n=17) compared to naïve-single (yellow circles, n=17) and mated (red squares, n=16, 5 heads/sample). Metabolites of the kynurenine pathways are highlighted in orange; serotonin is highlighted in yellow. NS *p*>0.05, \**p*<0.05, \*\**p*<0.01, \*\*\**p*<0.001. Statistical analysis was performed by ANOVA or Kruskal-Wallis with post-hoc Tukey’s or Friedman test.

**Figure S6.**
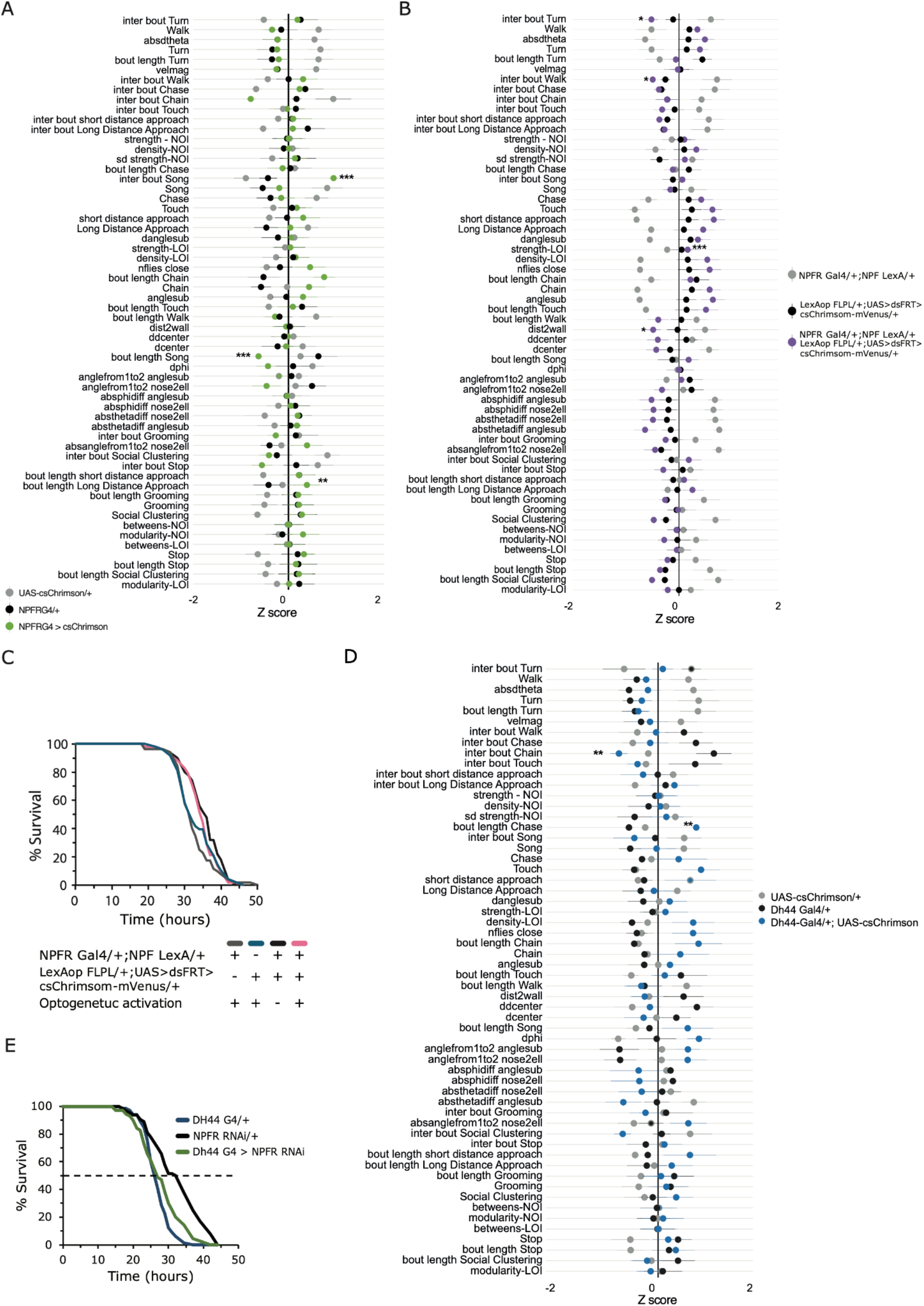
**A.** Behavioral signatures of male-male social interaction within the FlyBowl system during the optogenetic activation of all NPFR neurons. Data is represented as normalized Z scores of 60 behavioral parameters. NPFRG4/+;UAS-csChrimson/+ males (green, n=14) and their genetic controls NPFR G4/+, UAS-csChrimson/+ (black and grey, respectively, n=10 each). ANOVA or Kruskal-Wallis with post-hoc Tukey’s or Dunn’s test with FDR correction for multiple comparisons was performed. **B**. Behavioral signatures of male-male social interaction within the FlyBowl system during the optogenetic activation of all NPFR-NPF neurons. Data is represented as normalized Z scores of 60 behavioral parameters. (purple, n=22) and their genetic controls NPFR G4/+;NPF LexA/+ (grey, n=22), LexAop-FLPL/+;UAS<dsFRT>cs-Chrimson-mVenus/+, (black, n=21). *p*>0.05 Kruskal-Wallis. ANOVA or Kruskal-Wallis with post-hoc Tukey’s or Dunn’s test, and FDR correction for multiple comparisons were performed. **C.** Starvation resistance assayed on NPFR^NPF^ neurons by crossing NPFRG4;+, NPF-LexA;+, +;LexAop-FlpL, +;UAS<dsFRT>cs-Chrimson-mVenus flies. Naïve experimental males (Light, pink), n=52 and their genetic controls (NPFR G4/+;NPF LexA/+, grey, n=52; LexAop-FLPL/+;UAS<dsFRT>cs-Chrimson-mVenus/+, dark blue, n=58) were exposed to red light three times a day for two days. NPFR-NPF flies that were not exposed to light served as a third control (Dark, black, n=50). Experimental flies (pink) did not exhibit significantly different resistance to starvation compared to control flies (gray) *p*>0.05. **D.** Behavioral signatures of male-male social interaction within the FlyBowl system during the optogenetic activation of Dh44 neurons. Data is represented as normalized Z scores of 60 behavioral parameters. n=13 for NPFR^TK^(red). n=13 TK-LexA;LexAop-FLPL (black), and n=12 NPFRG4;UAS<dsFRT>csChrimson-mVenus (gray). \**p*<0.05, \*\**p*<0.01, \*\*\**p*<0.001. ANOVA or Kruskal-Wallis with post-hoc Tukey’s or Dunn’s test, and FDR correction for multiple comparisons were performed. **E.** Starvation resistance assayed on Dh44 G4/NPFR RNAi (green, n=70) flies and their genetic controls Dh44 G4/+ (blue, n=66) and NPFR RNAi/+ (black, n=64). No significant difference was observed among experimental flies and the controls, *p*>0.05. Pairwise log-rank test with FDR correction for multiple comparisons was performed.

**Figure S7.**
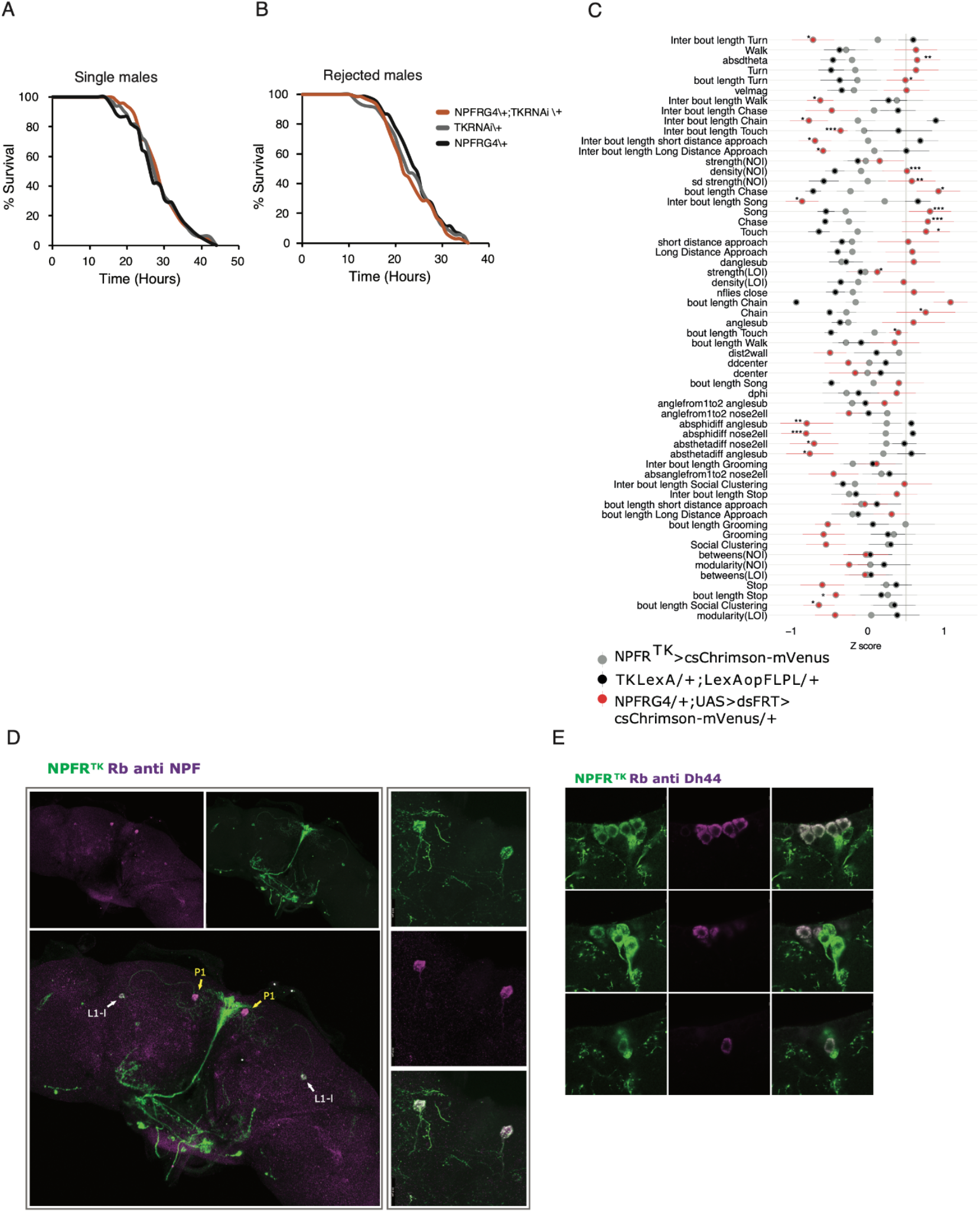
**A.** Knock down of tk in NPFR neurons does not affect sensitivity to starvation of naïve-single. Experimental single housed NPFRG4/+;tkRNAi/+ (orange, n=50) and the genetic controls TK RNAi/+ (gray, n=33) and NPFR G4/+ (black, n=45). **B.** NPFRG4/+;tkRNAi/+ males and their genetic controls were subjected to rejection and their resistance to starvation was assayed. No significant difference in resistance to starvation in NPFR G4;tk RNAi flies (orange, n=79) compared to genetic controls (gray, n=90 and black, n=70) was observed. Pairwise log-rank test with FDR correction for multiple comparisons was performed for A,B. **C.** Behavioral signatures of male-male social interaction within the FlyBowl system during the optogenetic activation of NPFR-TK neurons. Data is represented as normalized Z scores of 60 behavioral parameters. NPFRG4/+;UAS-csChrimson/+ males (green, n=14) and their genetic controls NPFR G4/+, UAS-csChrimson/+ (black and grey, respectively, n=10 each). ANOVA or Kruskal-Wallis with post-hoc Tukey’s or Dunn’s test with FDR correction for multiple comparisons was performed. **D.** Six NPFR^TK^ (green) neurons colocalize with DH44 (magenta, endogenous Dh44 expression). **E.** Right: colocalization of NPFR^TK^ neurons (green) and NPF+ neurons (magenta, endogenous NPF expression), indicated by arrows. White arrows indicate L1-l neurons, yellow arrows indicate P1 neurons. **Left:** A closeup to two NPFR^TK^ NPF+ neurons (P1).

## Notes

### Competing Interest Statement

The authors have declared no competing interest.

### Summary of Updates

We performed new experiments based on comments

